# The FMRF-NH_2_ Gated Sodium Channel of *Biomphalaria glabrata*: Localization and Expression Following Infection by *Schistosoma mansoni*

**DOI:** 10.1101/2022.12.15.520648

**Authors:** Laura Vicente-Rodríguez, Amanda Torres, Anthony Hernández-Vázquez, Mariela Rosa-Casillas, Dina P. Bracho-Rincón, Paola Méndez de Jesús, Martine Behra, Joshua J.C. Rosenthal, Mark W. Miller

## Abstract

The neglected tropical disease schistosomiasis impacts the lives of over 700 million people globally. *Schistosoma mansoni*, the trematode parasite that causes the most common type of schistosomiasis, requires planorbid pond snails of the genus *Biomphalaria* to support its larval development and transformation to the form that can infect humans. A greater understanding of neural signaling systems that are specific to the *Biomphalaria* intermediate host could lead to novel strategies for parasite or snail control. This study characterized a *Biomphalaria glabrata* neural receptor that is gated by the molluscan neuropeptide FMRF-NH_2_. The *Biomphalaria glabrata* FMRF-NH_2_ gated sodium channel (*Bgl*-FaNaC) amino acid sequence was highly conserved with FaNaCs found in related gastropods, especially the planorbid *Planorbella trivolvis* (91% sequence identity). In common with the *P. trivolvis* FaNaC, the *B. glabrata* receptor exhibited a low affinity (EC_50_: 3 × 10^−4^ M) and high specificity for the FMRF-NH_2_ agonist. Its expression in the central nervous system, detected by immunohistochemistry and *in situ* hybridization, was widespread, with the protein localized mainly to neuronal fibers and the mRNA confined to cell bodies. Colocalization was observed with the FMRF-NH_2_ tetrapeptide precursor in some neurons associated with male mating behavior. At the mRNA level, *Bgl*-FaNaC expression in the visceral and left parietal ganglia decreased at 20 days post infection by *S. mansoni* and in the shedding phase. Altered FMRF-NH_2_ signaling could be vital for parasite survival and proliferation in its snail intermediate host.

## 1. Introduction

Pond snails of the genus *Biomphalaria* serve as intermediate hosts for *Schistosoma mansoni*, the causative agent for the most widespread form of intestinal schistosomiasis (Maldonado and Perkins 1967; Wright 1971). Within their snail hosts, larval trematodes multiply and transform into the cercarial form that can infect humans (Rollinson and Chappell 2002; Toledo and Fried 2011). Strategies for elimination of schistosomiasis include increased sanitation, large-scale preventive chemotherapy, and snail control (Bergquist and Gray 2019; Allan et al. 2020; WHO Fact Sheets 2022).

Neuropeptide signaling systems are promising molecular targets for pesticide and parasiticide drug development (Maule et al. 2002; Geary and Maule 2010; McVeigh et al. 2011). In contrast to the classical neurotransmitter systems that are presently common targets for pest control, some neuropeptides and their receptors are limited to specific invertebrate clades, reducing concerns of widespread toxicity (Mousley et al. 2005; Martin and Robertson 2010; McVeigh et al. 2011). The FMRF-NH_2_ family of neuropeptides holds potential for drug development due to its pervasive role in the behavior and neuromuscular physiology of major arthropod and helminth parasites and pests (Mousley et al. 2004; McVeigh et al. 2005; 2009). While FMRF-NH_2_ was initially purified from molluscs (Price and Greenberg 1977) and has been intensively studied in several gastropod species (Cottrell 1993; Santama and Benjamin 2000; Zatylny-Gaudin and Favrel 2014), the potential utility of this peptide signaling system for snail control interventions has received less scrutiny.

In common with most neuropeptides, the gastropod FMRF-NH_2_-related peptides activate G-protein coupled receptors (GPCRs) that regulate ion channels via second messenger systems (Higgins et al. 1978; Piomelli et al. 1987; Willoughby et al. 1999a, b; Phan et al. 2022). In addition, FMRF-NH_2_ directly activates a member of the degenerin / epithelial sodium channel (DEG/ENaC) superfamily of channels that do not require second messenger signaling (Green et al. 1994; Lingueglia et al. 1995, 2006; Cottrell 1997). Ionotropic DEG/ENaC channels are expressed in distinct cell types and tissues and are gated by varied stimuli, including mechanical forces and protons (Benos and Stanton 1999; Kellenberger and Schild 2002; Eastwood and Goodman 2012). In the nervous system, members of this receptor family participate in a range of functions, including mechanotransduction, nociception, and synaptic plasticity (Waldman and Lazdunski 1997; O’hagan et al. 2005; Bianchi 2007). The role of the FMRF-NH_2_ activated sodium channel (FaNaC) in the neural circuits that regulate gastropod physiology and behavior remains unspecified.

Recently, we examined the precursor organization of the *B. glabrata* FMRF-NH_2_-related neuropeptides and localized their expression in the CNS (Rolón-Martínez et al. 2021). As our understanding of this neuropeptide system will be broadened by defining its complementary receptors, the present study characterized the *B. glabrata* FMRF-NH_2_ gated sodium channel, localized its expression in the CNS, and explored whether its expression may be modified following exposure to *Schistosoma mansoni*.

## 2 Materials and Methods

### 2.1 Specimens

*B. glabrata* snails were bred in the animal facility of the Institute of Neurobiology, University of Puerto Rico Medical Sciences Campus. Snails were maintained in aquaria at room temperature under a 12-hour light/dark cycle. They were fed with lettuce *ad libitum*. All protocols were approved by the Institutional Animal Care and Use Committee (IACUC) of the University of Puerto Rico Medical Sciences Campus (Protocol #3220110).

*B. glabrata* exposure to *S. mansoni* miracidia was conducted at the Biomedical Research Institute (BRI, Rockville MD). Schistosome eggs were obtained from mouse livers and hatched using the BRI protocols (Lewis et al. 1986). Snails were incubated with miracidia (target: 5 per snail) for two hours. Tissues from infected specimens were dissected and collected at the BRI at 20 (prepatent) and 35 (shedding) days post-exposure. Shedding was induced by exposure of snails to direct light. All snails used for this study ranged from 8–15 mm in shell diameter.

### 2.2 Electrophysiology

The cDNA encoding *Bgl-*FaNaC sequence was codon-optimized for *Xenopus laevis* and synthesized by Integrated DNA Technology Co. (IDT, Coraville IA). The Flag-tag protein affinity sequence (DYKDDDDK; Sigma-Aldrich) was added to the N-terminal to facilitate confirmation of expression. The cDNA was subcloned into the *Xenopus* expression vector pGEM HE as described previously (Ferrick-Kiddie et al. 2017). The *Bgl*-FaNaC full-length RNA was transcribed *in vitro*, capped and polyadenylated using the T7 mScript Standard mRNA Production System (CellScript).

All animal care and experimental procedures involving *Xenopus* were approved by the University of Puerto Rico Institutional Animal Care and Use Committee and were performed following relevant guidelines and regulations. Oocytes were obtained from dissected ovaries from adult specimens of *Xenopus laevis* (Xenopus Express, Brooksville FL). They were dispersed with type II collagenase and manually defolliculated. RNA injections were performed in pre-selected oocytes from stages V and VI. Oocytes were injected with 38.6 nL of *Bgl*-FaNaC encoding RNA at a concentration of 20 ng/uL using a Nanoliter 2000 microinjector (World Precision Instruments). Maximum expression levels were observed 6 days following injection.

Injected oocytes were placed at the bottom of a plastic recording chamber (3 ml volume) lined with a nylon mesh and continuously perfused with ND96 (96 mM NaCl, 2 mM KCl, 1.8 mM CaCl_2_, 1 mM MgCl_2_, 5 mM HEPES, pH, 7.6; see detailed methods in Ferrick-Kiddie et al. 2017). An OC725B oocyte clamp (Warner Instruments LLC., Hamden CT) was used to clamp the membrane potential at −60 mV with independent recording (0.1 M potassium chloride) and current passing (3 M potassium chloride) microelectrodes. Data were acquired by Digidata 1200 and the Clampex (V.6, Axon Instruments, Union City CA) and analyzed using AxoScope 10.7. All experiments were carried out at room temperature (20-25 °C).

Synthetic FMRF-NH_2_, FLRF-NH_2_, FIRF-NH_2_, pQFYRI-NH_2_, and GDPFLRF-NH_2_ were purchased from GL Biochem (Shanghai, China). All peptides were dissolved in ND96 in a stock of 1 mM. Peptide stocks were frozen at −20 °C and serial dilutions were freshly prepared before each experiment. Peptides were applied manually using a pipettor (1 mL, upstream).

### 2.3 Antibody Validation

Affinity purified rabbit polyclonal antibodies were generated against the amino terminus intracellular domain of the *B. glabrata* FMRF-NH_2_-gated sodium channel (KYTSPDAKPSMSTS-C; residues 2-15, Fig. 1a, 2) by GL Biochem Ltd., Shanghai, China (ELISA titer > 1:128,000). Solid phase specificity was confirmed with dot blots of serial antigen dilutions (2 μL) applied to nitrocellulose membrane (Bio-Rad 0.45 µm; Fig. 1b). Membranes were allowed to air dry (1 h) and then incubated with blocking buffer (1 h, room temperature, shaking). Membranes were then incubated overnight with the anti-FaNaC primary antibody diluted 1:200 in blocking solution (4 °C, shaking). For preabsorption controls, the antibody was pre-incubated with the peptide antigen (10^−4^ M) overnight prior to application to the membrane. The membranes were then washed three times for 15 minutes and incubated with goat anti-rabbit IgG (H+L) second antibody conjugated to HRP (1:4000 in block solution, 1 h, room temperature). They were then washed 4 × 15 min in Tris-buffered saline, 0.01% Tween 20 (TBS-T), and then transferred to SuperSignal™ West Femto Maximum Sensitivity Substrate. The enzyme-substrate reaction was allowed to proceed for five minutes before visualization.

**Fig. 1.**
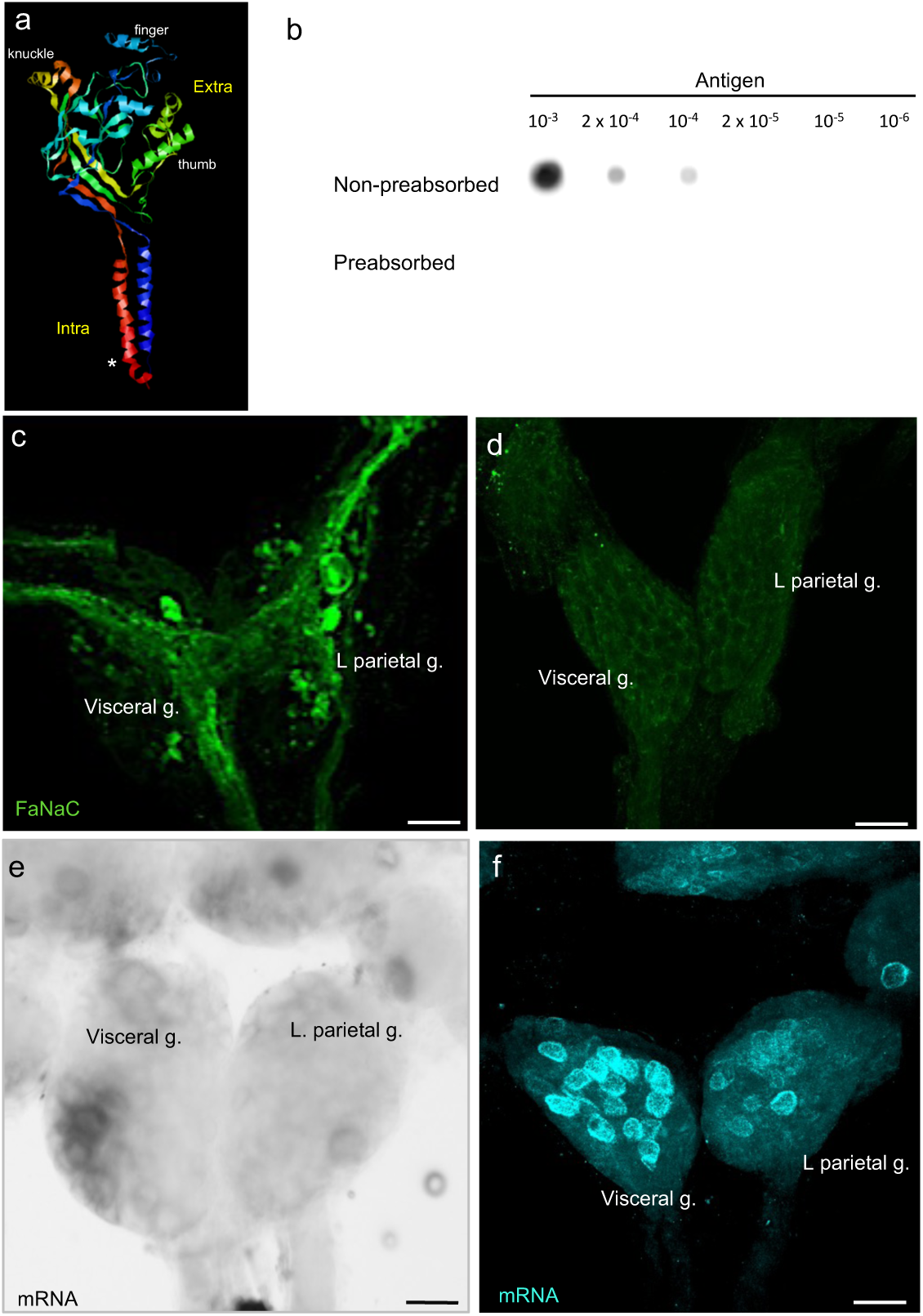
Localization of *Bgl*-FaNaC expression. **a:** Ribbon model of *Bgl*-FaNaC channel subunit produced with the Open RasMol Molecular Graphics Visualization Tool (v2.7; www.penrasmol.org). The channel comprises two intracellular (*Intra*) N- and C-termini and a large extracellular (*Extra*) domain. Features of the ‘clenched fist’ conformation are labeled (see Eastwood et al. 2012). The antibody used in this study was generated against a 14-residue domain near the amino terminus of the channel (*asterisk*). The selected sequence was based upon an analysis of antigenicity performed by GL Biochem (Shanghai, China). **b:** Dot blot controls demonstrate specificity of the *Bgl*-FaNaC antibody. Upper row: serial dilutions of a 2 µM antigen solution were blotted (2 µl) and probed with a 1:200 dilution of the antibody used in this study. Lower row: Preabsorption of the antibody with the antigen peptide (1 × 10^−4^ M, overnight) eliminated recognition of the blotted peptide. **c:** Wholemount immunolabeling of the ventral surface of the visceral and left parietal ganglia. Intense FaNaC labeling (*green*) was present in fiber systems coursing through the ganglia. **d:** FaNaC labeling was eliminated following antibody preabsorption with the antigen. **e:** The signal produced with the digoxygenin *in situ* hybridization protocol was confined to the cell bodies of neurons in the visceral and left parietal ganglia. **f:** Detection of FaNaC mRNA using the Hybridization Chain Reaction technique produced defined labeling (*cyan*) in the cell bodies of neurons in the visceral and left parietal ganglia. All calibration bars = 50 µm.

**Fig. 2.**
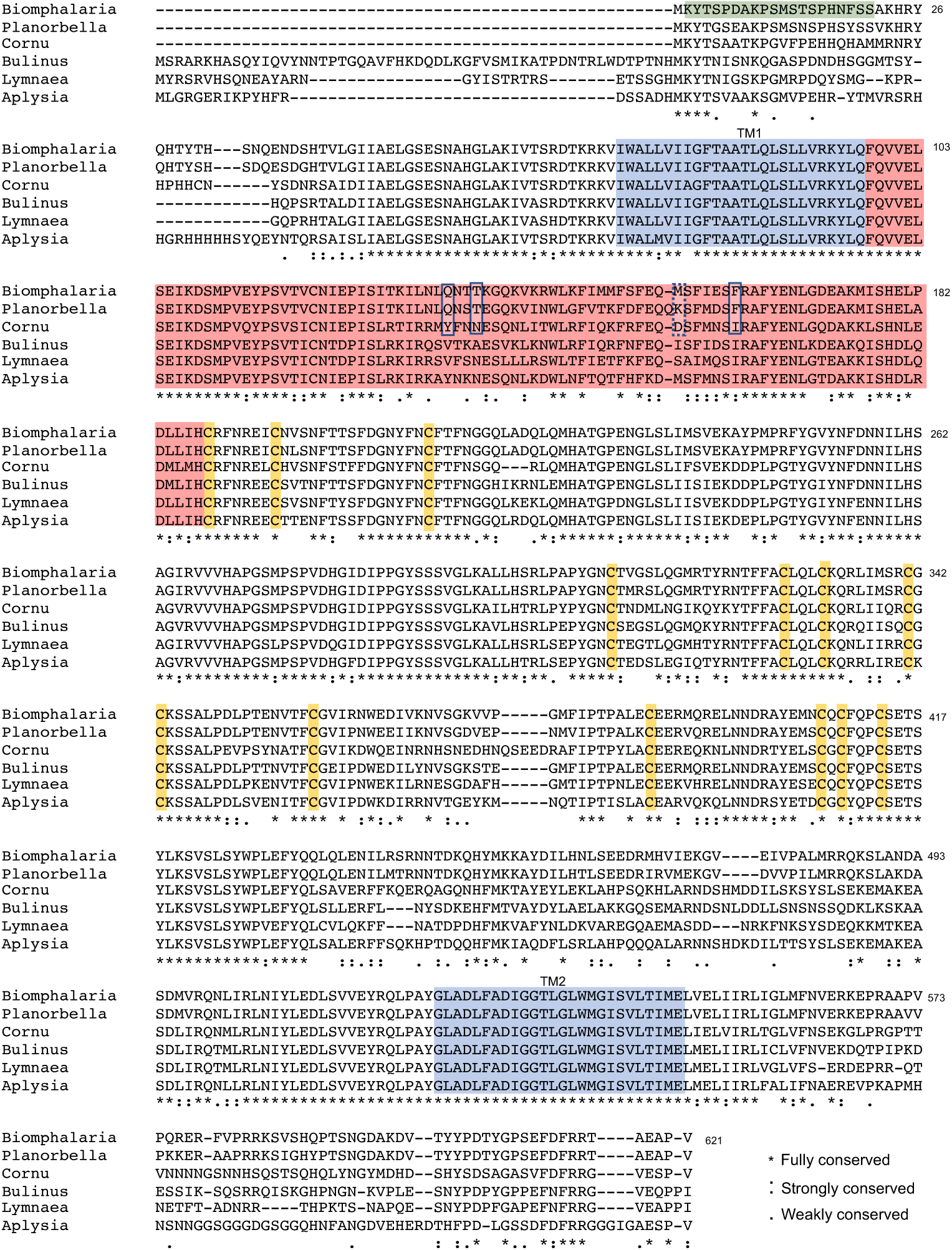
Sequence alignment of the *Bgl*-FaNaC with FaNaCs reported for other gastropods. Sequences include: *Planorbella trivolvis* (GenBank ID: AAF80601.1; Jeziorski et al. 2000), C*ornu aspersum*, (GenBank ID: CAA63084.1; Lingueglia et al. 1995), *Bulinus truncatus* (GenBank ID: KAH9494543.1), *Lymnaea stagnalis* (GenBank ID: AAK20896.1; Perry et al. 2001), and *Aplysia kurodai* (GenBank ID: AB206707.1). Amino acid numbering corresponds to *Bgl*-FaNaC. Sequence of the peptide used to generate the antibody used in this study is shaded light green. Orange shading highlights conserved cysteine residues that are thought to contribute to disulfide bridges in the finger region of the channel. Light blue shading denotes the two highly conserved transmembrane domains, TM1 and TM2. The less conserved region that is proposed to account for peptide recognition is shaded light red. Within this domain, four specific residues that influence the concentration-response relationship are outlined (see Niu et al. 2016). The alignment was generated with the T-Coffee web server (Notredame et al. 2000).

Preabsorption experiments on fixed *Biomphalaria* nervous tissue also verified the specificity of antigen detection. A 1:200 dilution of the anti-FaNaC antibody produced strong immunofluorescence (Fig. 1c) that was eliminated when the antibody was preabsorbed with the antigen prior to tissue incubation (5 × 10^−4^ M, overnight; Fig. 1d). Together, the preabsorption experiments supported the sensitivity and specificity of the antibody used in this study. They also provided guidance for primary and secondary antibody dilutions to use for antigen detection. Signals were eliminated when primary antibody incubation was omitted from the protocol (not shown).

### 2.4 Whole-mount immunohistochemistry

Standard wholemount immunohistochemical protocols were followed (Habib et al. 2015; Vaasjo et al. 2018). Tissues were dissected in normal saline (in mmol l^−1^: NaCl 51.3, KCl 1.7, MgCl_2_ 1.5, CaCl_2_ 4.1, HEPES 5, pH 7.8.), and pinned in a Sylgard plate. Ganglia were incubated in protease (0.5%; Type XIV, Sigma #P5147); 7-10 min), washed thoroughly with normal saline, and then fixed in 4% paraformaldehyde (1 h, room temperature).

Fixed tissues were washed 5 × 20 min in PTA (0.1 M phosphate buffer containing 2% Triton X-100 and 0.1% sodium azide) at room temperature. Following pre-incubation with normal goat serum (0.8%, 3-12 h, room temperature), tissues were transferred to the primary antibody (rabbit polyclonal raised against KYTSPDAKPSMSTS-C (GL Biochem, Shanghai, China; 1:200 dilution in PTA, 3-5 days). Samples were washed (5 × 20 min in PTA) and incubated in second antibodies conjugated to a fluorescent marker (Alexa 488 goat anti-rabbit IgG (H+L) conjugate; Molecular Probes, Eugene OR) at dilutions ranging from 1:500 to 1: 1,000). Quality of the staining was assessed on a Nikon Eclipse fluorescence microscope prior to imaging. Confocal imaging was performed on a Nikon A1R Confocal Laser Microscope using the NIS Elements AR software package (Version. 4.5, Nikon Instruments). Stacks, z-series, overlays, and calibrations were produced using the Fiji software (v. 2.00, NIH public domain).

### 2.5 In situ hybridization

#### Whole-mount in situ hybridization with chromogenic detection

Digoxygenin (DIG)-labeled probes were produced with the SuperScript^TM^ III One-Step RT-PCR kit (Sigma 12574-026) using specific primers for the *Bgl-*FaNaC transcript (Forward: CCAGCATGTCTACCTCACCGCAC, Reverse: CTCCGTAGGCAAGTCCGGCAAGG). T7 and T3 promoter sequences were appended to the forward and reverse primers, respectively. The expected amplicon length was 1,034 bp. Ganglia were dehydrated and rehydrated in methanol/10x PBS graded solutions (25%, 50%, and 75% methanol) for 5 min and then rinsed in PBST (1% Tween 20) for 5 min. Tissues were digested with 2 mg/ml of Proteinase K (Ambion) for 7 min, and fixed with 4% paraformaldehyde (room temperature, 45 min). After rinsing 5 × 5 min in blocking buffer (1 × PBS, 0.1% Tween 20, 0.1% BSA, 1% DMSO), ganglia were pre-hybridized in hybridization mix (50% formamide, 5 × SSC, 1 mg / ml yeast RNA, μm/ml heparin, 0.1% Tween 20, 5 mM EDTA, 9 mM citric acid, in DEPC treated water) for 4-6 h. Hybridization was performed with the pre-heated antisense probe (final concentration = 1 ng/μl) overnight at 65 °C. Tissues were washed with a graded series of hybridization mix solutions (75%, 50%, 25%) in 2x SSC for 10 min each, followed by two 30 min washes with 0.2x SSC, all at 65 °C. Tissues were then rinsed in 0.2x SSC graded solutions (75%, 50%, and 25%) in PBST for 10 min each at RT. For detection, specimens were pre-incubated in blocking buffer for 4-6 h and then incubated in anti-DIG antibody conjugated to alkaline phosphatase (1/3000 blocking buffer, overnight). Finally, tissues were rinsed in PBST 6 × 15 min followed by two 5 min washes in alkaline phosphatase buffer (100 mM Tris pH 9.5, 50 mM MgCl_2_, 100 mM NaCl, 0.1% Tween 20, levamisole). Signal development was performed with 100% BM-Purple (Roche) in the dark (Fig. 1e).

### 2.6 Hybridization Chain Reaction (HCR) fluorescence *in situ* hybridization

HCR RNA-FISH methods were adapted from the Molecular Instruments, Inc. (Los Angeles, CA) protocols website (https://www.molecularinstruments.com/). The *B. glabrata* CNS was dissected in normal saline (mmol l^−1^: NaCl 51.3, KCl 1.7, MgCl_2_ 1.5, CaCl_2_ 4.1, HEPES 5, pH 7.8) and pinned on Sylgard-lined plates. Tissues were exposed to protease (0.5%; Type XIV, Sigma) diluted in normal saline for 7-10 minutes and fixed in 4% paraformaldehyde overnight. The CNS was washed 5 times for 15 minutes each with PTwA (0.1 M phosphate buffer containing 2% Tween 20 and 0.1% sodium azide). Samples were then pre-hybridized in hybridization buffer (Molecular Instruments, Inc.) for 30 minutes, and then hybridized overnight at 37 ᵒC with a probe set generated for the *Bgl*-FaNaC (FaNaC/ LOT PRI 987) transcript or a cocktail of probe sets that also included the *Bgl*-FaRP1 precursor (LOT PRI087) and the *Bgl*-FaRP2 precursor (LOT PRI298). Probes were diluted in hybridization buffer at a final concentration of 4 nmol/µL each. Samples were washed 4 × 15 minutes with wash buffer (Molecular Instruments, Inc.) and incubated at room temperature with amplification buffer (Molecular Instruments, Inc.) for 30 minutes. Meanwhile, aliquots of the hair pin amplifiers (h1 and h2; 5 µL each, 100 µM) were heated at 95 ᵒC for 90 s and then placed in a dark box for 30 minutes at room temperature. Following cooling, 5 µL of each hair pin was added to 250 µL of amplification solution. Tissues were transferred to the amplification solution and incubated overnight at room temperature in a dark box. The following day, samples were washed 5 times (10 minutes each) with 5x SSCT at room temperature. Results were visualized on a Nikon Eclipse epi-fluorescent microscope prior to confocal imaging on a Nikon A1R Confocal Laser Microscope using the NIS Elements AR software package (Fig. 1f).

### 2.7 Analysis of expression

As the fluorescence mRNA detection provided superior clarity and definition (Fig. 1e, f), all expression measurements were obtained from samples using the HCR method. Images from control and infected samples were obtained using the same settings on the NIS Elements data acquisition program. The Nd2 files were opened in Fiji 4.5 and the number of labeled neurons on the dorsal and ventral surfaces of each ganglion were counted manually by an experimenter blinded to the treatment. The mean gray value cut-off for positive expression was set at 15. Neurons with a diameter less than 10 µm were excluded. Overall mean gray values were obtained from a region of interest (ROI) demarcated with the Fiji “free hand selection” tool. For *Bgl*-FaRP1 and *Bgl*-FaRP2 expression analysis, specific clusters (B group, F group and E group; see Rolón-Martínez et al., 2021) were selected as the ROI for gray value measurements. As *Bgl*-FaNaC expression was more dispersed, the perimeter of each ganglion was traced to demarcate the ROI.

### 2.8 Statistical analysis

Data are presented as mean ± standard error of the mean (SEM). The number of specimens used for each measurement is included in the graphical representation and indicated in the figure legends. Statistical significance was determined using the Brown-Forsythe and Welch one-way analysis of variances (ANOVA) test with Dunnett’s multiple comparison as follow-up test. Tests were performed and graphs were generated using GraphPad Prism 9.2.0.

## 3 Results

### 3.1 The *B. glabrata* FaNaC structure and function

A transcriptome generated from twelve pooled *B. glabrata* central nervous systems (Rolón-Martínez et al. 2021) yielded a transcript with homology to the FMRF-NH_2_ gated sodium channel of *Cornu aspersum* (*Helix aspersa*). This 4444 nucleotide sequence (GenBank Accession number OP066530) encompassed a 1395 nucleotide 5’ untranslated sequence, an open reading frame (ORF) encoding a 621 amino acid protein termed *Bgl*-FaNaC, and a 1186 nucleotide 3’ untranslated sequence. The *Bg*FaNaC amino acid sequence was identical to the *B. glabrata* FMRFamide-activated Amiloride-sensitive Sodium Channel-like Protein previously derived from genomic sequence (Accession number XP_013063507).

Alignment of the *Bgl*-FaNaC amino acid sequence with gastropod FaNaCs reported previously confirmed significant sequence identity (Fig. 2; *Planorbella trivolvis*: 91%, *Bulinus truncatus*: 70%, *Lymnaea stagnalis*: 70%, *Aplysia californica*: 67%, *Cornu aspersum*: 66%). A ribbon model generated with the Open RasMol Molecular Graphics Visualization Tool (Fig. 1a) retained several conserved characteristics of the DEG/ENaC ion channel superfamily, including two highly conserved transmembrane domains (Fig. 2, shaded blue), intracellular N- and C-termini, and a large “clenched fist” ectodomain. Thirteen cysteine residues located in the finger region were fully conserved (Fig. 2, orange shading). Lower conservation was observed in the thumb region, which is thought to confer agonist specificity and efficacy of FaNaC receptors (Cottrell et al. 2001; Cottrell 2005; Niu et al. 2016)

The *Bgl*-FaNaC coding sequence was optimized for expression in *Xenopus*, cloned into an expression vector, and injected into oocytes. Bath application of FMRF-NH_2_ (7.5 × 10^−4^ M; 1 mL, upstream) produced an inward current (Fig. 3b). No currents followed application of related peptides that are encoded on the *Biomphalaria* FMRF-NH_2_ precursors (Rolón-Martínez et al., 2021), including FLRF-NH_2_, (Fig. 3b), pQFYRI-NH_2_ (Fig. 2c), and FIRF-NH_2_ (Fig. 3c), and the heptapeptide GDPFLRF-NH_2_ (Fig. 3d). The FMRF-NH_2_ responses were concentration-dependent, with a mean EC_50_ of 3.3 × 10^−4^ M (Fig. 4). Desensitization was not observed under the conditions of peptide delivery (Lingueglia et al. 1995; Jeziorski et al. 2000; Schanuel et al. 2008).

**Fig. 3.**
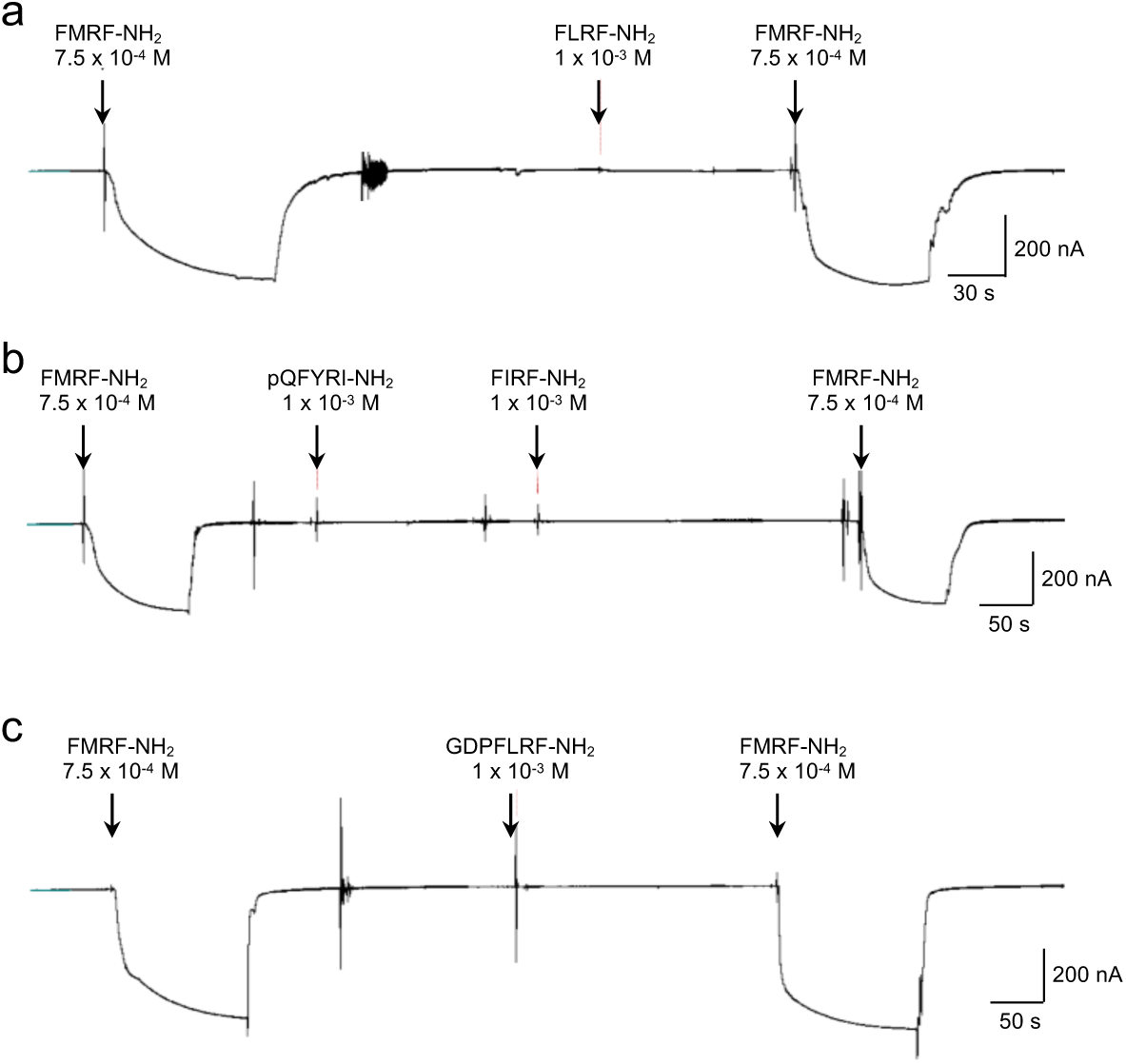
*Bgl*-FaNaC specificity demonstrated with heterologous expression in *Xenopus* oocytes. Currents were recorded with a two-electrode voltage clamp configuration. **a:** Application of FMRF-NH_2_ (7.5 × 10^−4^ M, 1 mL) produced large inward currents. No response was detected following application of FLRF-NH_2_ (1 mM). **b:** Tests with two peptides encoded on the FMRF-NH_2_ tetrapeptide precursor *Bgl-FaRP1* (Rolón-Martínez et al. 2021), pQFYRI-NH_2_ (1 mM) and FIRF-NH_2_ (1 mM), substantiated the specificity of the *Bg*FaNaC. **c:** No response was elicited by GDPFLRF-NH_2_, a product of the heptapeptide precursor *Bgl*-FaRP2 produced by alternative splicing of the FMRF-NH_2_ message.

**Fig. 4.**
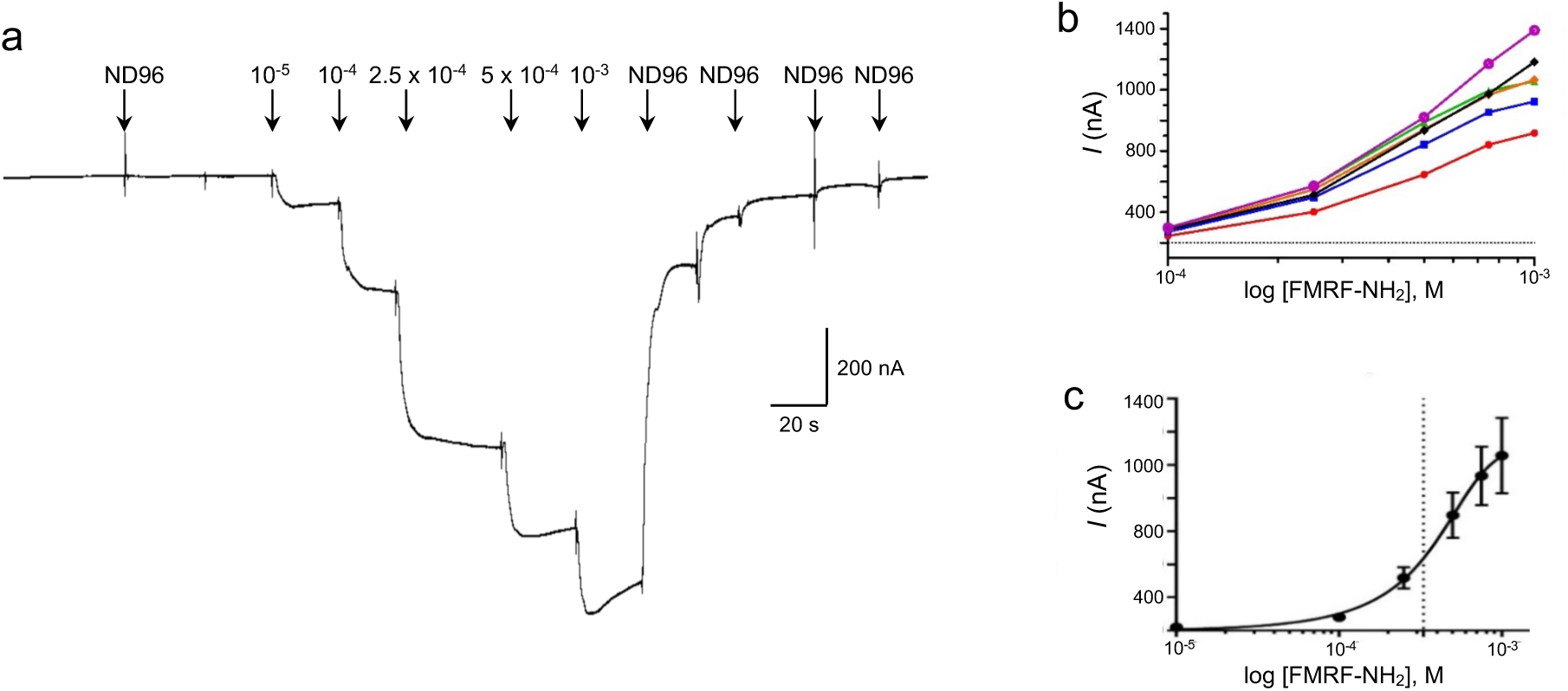
Concentration dependence of *Bgl*-FaNaC responses to FMRF-NH_2_. **a:** Application of graded concentrations of FMRF-NH_2_ (10^−5^ M – 10^−3^ M; 1 mL) produced increased inward currents in the *Xenopus* heterologous expression system. **b:** Uniform concentration-response profiles were obtained from five oocytes (color coded). **c:** Averaged data produced an EC_50_ of approximately 3 × 10^−4^ M (dashed line).

### 3.2 *Bgl*-FaNaC localization

A polyclonal rabbit antibody was generated against residues 2-15 of the *B. glabrata* FaNaC (Fig. 1a, 2). Immunohistochemical processing of wholemount central nervous systems labeled a widespread network, with cell bodies located in all ganglia, and abundant fiber systems within the peripheral nerves (5, 6). Dense fiber tracts also coursed through the central neuropil of the ganglia, often obscuring underlying cell bodies (Fig. 1c, 5a, c, d).

**Fig. 5.**
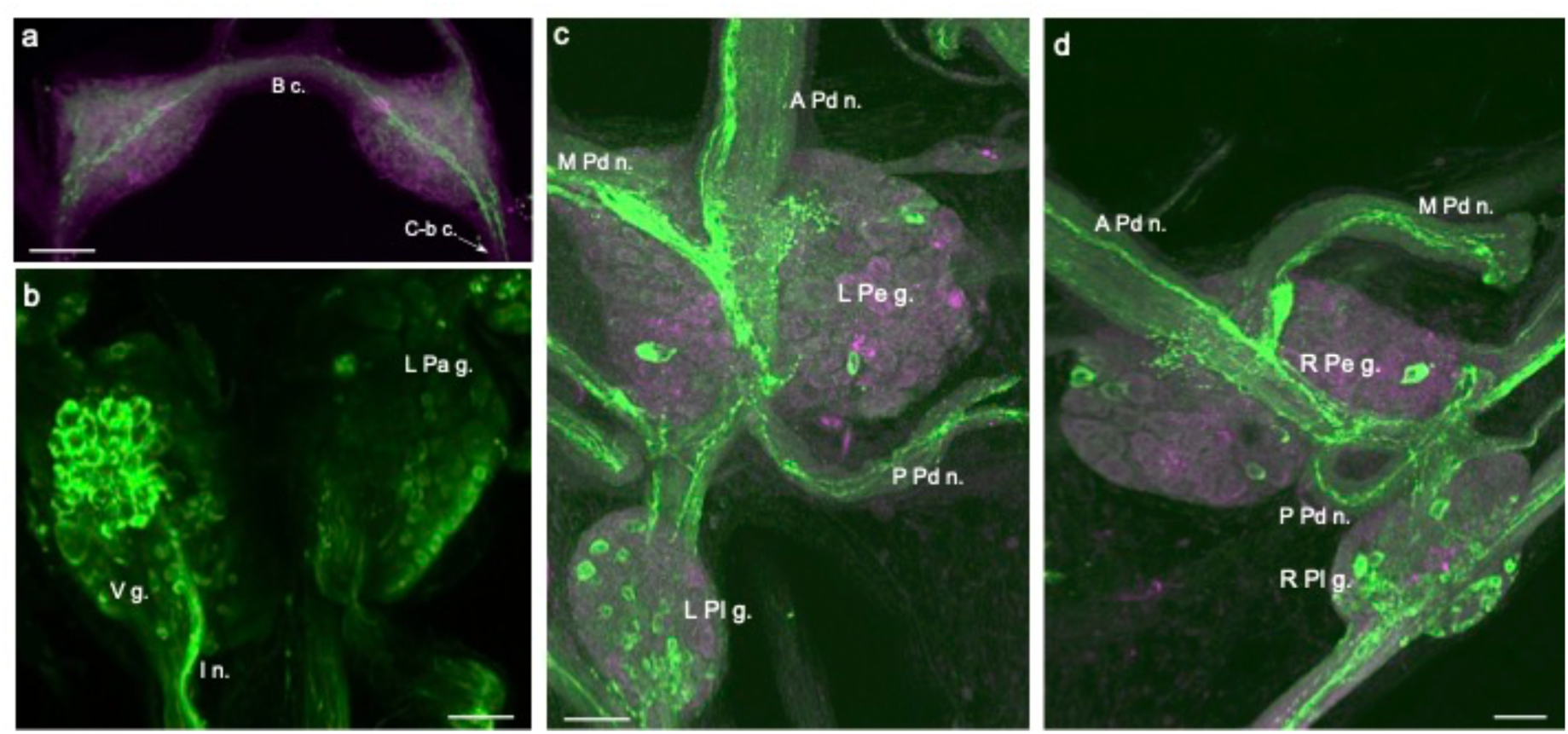
FaNaC-like immunoreactivity in the CNS of *Biomphalaria*. **a:** FaNaC-li fibers originating from the cerebral-buccal connective (*C-b c.*) crossed the buccal commissure (*B c.*). They coursed through each buccal hemiganglion giving rise to a diffuse network that permeated the central neuropil. *Calibration bar* = 50 µm. **b:** Intense labeling was observed in a cluster of cell bodies on the ventral surface of the visceral ganglion (*V g.*). These cells appeared to give rise to a compact bundle of fibers in the intestinal nerve (*I n.*). *Calibration bar* = 50 µm. **c:** FaNaC-li fibers coursed through the center of the left pedal ganglion (*L Pe g.*), projecting into the anterior, medial, and posterior pedal nerves (*A Pd n.*, *M Pd n.*, *P Pd n.*). Dispersed small neurons were located in the left pleural ganglion (*L Pl g.*). Dorsal surface shown. *Calibration bar* = 50 µm. **d:** FaNaC-li fibers coursed through the center of the right pedal ganglion (*R Pe g.*), projecting into the anterior, medial, and posterior pedal nerves (*A Pd n.*, *M Pd n.*, *P Pd n.*). Small neurons were located in the right pleural ganglion (*R Pl g.*). Dorsal surface shown. *Calibration bar* = 50 µm.

In the pedal ganglia, immunohistochemical FaNaC labeling was detected in two identified giant neurons, the dopaminergic left pedal dorsal 1 (LPeD1, Fig. 6a, b; see Vallejo et al. 2014) and the serotonergic right pedal dorsal 1 (RPeD1, Fig. 6c, d; see Delgado et al. 2012). As reported previously in *Helisoma* (Davey et al. 2001), FaNaC labeling of the LPeD1 cell body appeared to be spatially aggregated (Fig. 6b). FaNaC labeling of the RPeD1 cell body was more uniform (Fig. 6d).

**Fig. 6.**
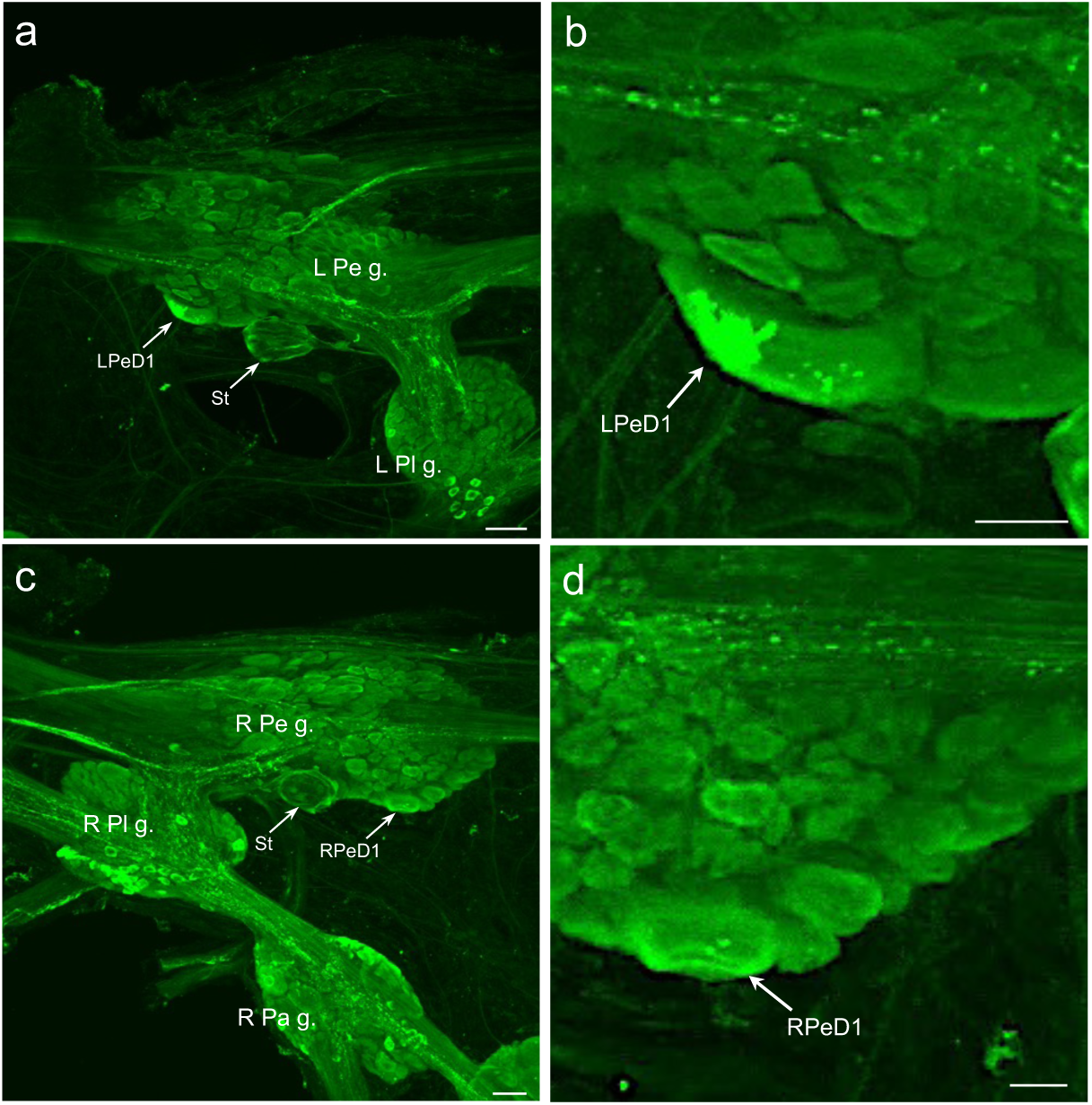
FaNaC-li in giant pedal ganglion neurons. **a:** The giant dopaminergic left pedal dorsal 1 (*LPeD1*) neuron was often optimally viewed from the ventral aspect of the ganglion (see Vallejo et al. 2014). FaNaC-li labeling was present in the LPeD1 and in the periphery of the statocyst (*St*). *Calibration bar* = 50 µm. **b:** Higher magnification revealed intense labeling of LPeD1 in a discrete region of the cell. *Calibration bar* = 20 µm. **c:** The giant right pedal dorsal 1 (*RPeD1*; Delgado et al. 2012) cell was apparent on the ventral aspect of the ganglion. The periphery of the right statocyst (*St*) was also labeled. *Calibration bar* = 50 µm. **d:** With higher magnification, FaNaC-li labeling of the RPeD1 was more distributed than in LPeD1. *Calibration bar* = 20 µm.

The two protocols used to detect *Bgl*-FaNaC mRNA in wholemount nervous systems yielded comparable results (see Materials and Methods). Chromogenic detection of cRNA probes (Fig 1c) and the fluorescent Hybridization Chain Reaction (HCR; Fig 1d) methods both produced labeling that was confined to cell somata, facilitating visualization of receptor expressing cells (Fig. 1e, f). *Bgl*-FaNaC expression, assessed with the HCR method, was widespread throughout the *B. glabrata* CNS (Fig. 10). All eleven ganglia contained neurons expressing this receptor. The left and right buccal ganglia had similar distributions of *Bgl*-FaNaC neurons. One prominent neuron was located on the superior dorsal surface each hemiganglion. This neuron was located medially, close to the buccal commissure (Fig. 7c). Around five medium sized neurons (20-25 µm) were located on the inferior margin and the lateral sides of each ganglion. In the cerebral ganglion, three prominent neurons expressing the receptor were present on the dorsomedial surface. Two giant neurons (25-40 µm) were positioned very close to the superior border and a cluster of smaller cells was located on the lateral side of each ganglion.

**Fig. 7.**
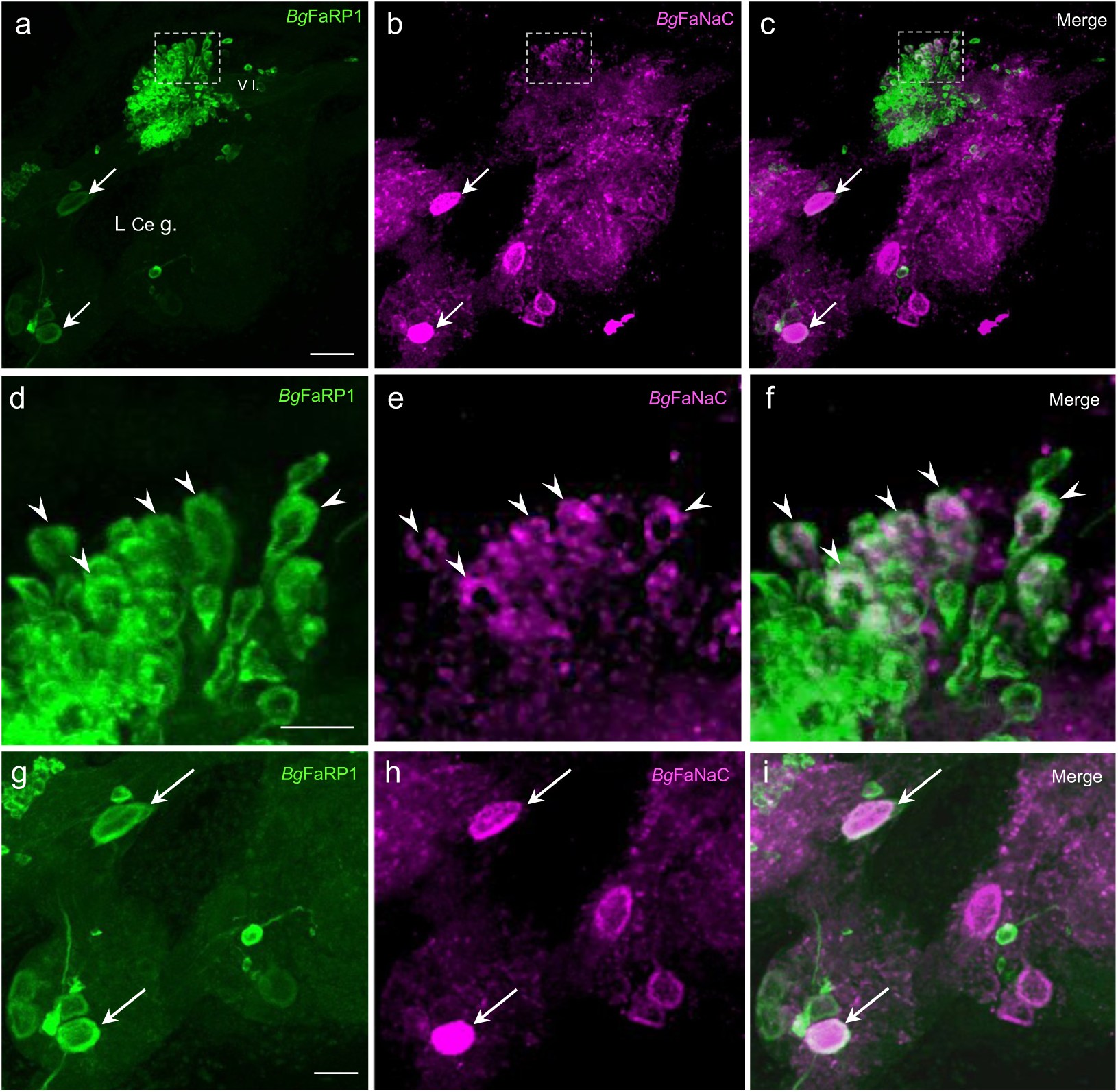
Colocalization of *Bgl*-FaNaC mRNA and *Bg*FaRP1 mRNA in neurons of the cerebral ganglion. Anterolateral quadrant of the left cerebral hemiganglion ventral surface shown. **a:** *Bgl*-FaRP1 mRNA was present in numerous small cells in the ventral lobe (*V l.*) of the left cerebral ganglion (*L Ce g.*; see Rolón-Martínez et al. 2021). *Calibration bar* = 30 µm, applies to **a-c**. **b:** A cluster of neurons in the lateral V l. express *Bgl*-FaNaC mRNA (*dashed rectangle*). Two larger neurons in the ventrolateral cerebral ganglion that express *Bgl*-FaRP1 also express *Bg*FaNaC (*arrows* in panels **a** and **b**). **c:** Overlay of panels **a** and **b** shows colocalization of *Bgl*-FaNaC mRNA and *Bgl-*FaRP1 mRNA in V l. cells. Neurons expressing both transcripts appear white. **d:** Region enclosed by *dashed rectangle* in **a** shown at higher magnification. Five neurons expressing *Bgl*-FaRP1 mRNA indicated by *arrowheads*. *Calibration bar* = 10 µm, applies to **d-f**. **e:** *Bgl*-FaNaC mRNA in the same field as **d**. Five labeled cells indicated by *arrowheads*. **f:** Overlay of panels **d** and **e** confirms co-expression of *Bgl*-FaRP1 and *Bgl*-FaNaC transcripts in a subset of neurons in the ventral lobe. **g:** Lower left quadrant of panel **a** shown at higher magnification. Two neurons expressing *Bgl*-FaRP1 mRNA indicated by *arrows*. *Calibration bar* = 20 µm, applies to **g-i**. **h:** *Bgl-*FaNaC mRNA in the same field as **g**. Two labeled cells indicated by *arrows*. **i:** Overlay of panels **g** and **h** confirms co-expression of *Bgl*-FaRP1 and *Bgl*-FaNaC transcripts in two anterolateral neurons extrinsic to the ventral lobe.

**Fig. 10.**
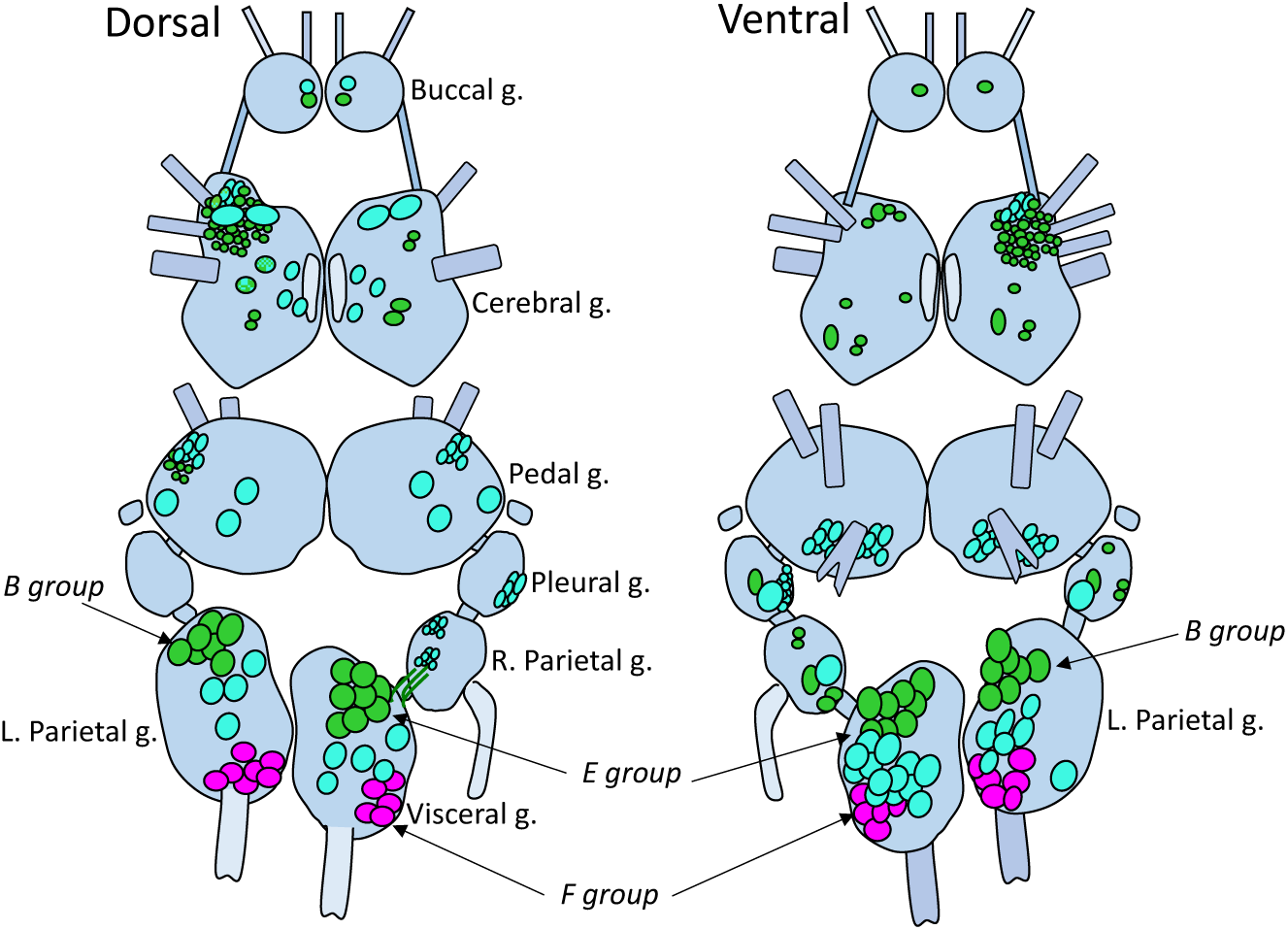
Schematic summary of expression patterns of the *B. glabrata* FaRP1 tetrapeptide precursor message (*green*), the FaRP2 heptapeptide precursor message (*magenta*), and the FaNaC message (*cyan*) in the *B glabrata* CNS.

The two pedal ganglia showed similar patterns of *Bgl*-FaNaC expression. Three prominent neurons were located on the dorsal surface. A giant neuron was located in the inferior and lateral border of each ganglion, a second prominent neuron was located in the center, and the third was located close to the inferior medial border. One cluster of approximately eight cells was present in the lateral and dorsal side of each ganglion. A cluster of roughly fifteen small neurons was located on the inferior border of the ventral side of each ganglion.

Both pleural ganglia contained one giant neuron (25-40 µm) expressing *Bgl*-FaNaC in the ventromedial and inferior border of each ganglion. The right parietal ganglion contained one giant neuron expressing *Bgl*-FaNaC in the ventromedial and inferior border of the ganglion. In the left parietal ganglion, one prominent positive neuron was present on the ventral surface in the inferior and lateral border, and another three to five prominent neurons were scattered through the ganglion (Fig. 14e, i). Twelve to fifteen medium sized (12-25 µm) positive neurons were located in the center and inferior lateral side of the ganglion. The visceral ganglion contained three to five prominent positive cells on the dorsal surface (Fig. 14e). A larger cluster of 12-15 medium sized neurons expressing *Bgl*-FaNaC was present on the ventral surface toward the lateral margin (Fig. 13a).

**Table I.** Number of neurons expressing Bgl-FaRP-1 and Bgl-FaRP-2 in visceral and left parietal ganglia 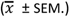.

|  |  | FaRP-1 |  |  |  | FaRP-2 |  |  |  |
| --- | --- | --- | --- | --- | --- | --- | --- | --- | --- |
|  |  | Control | 20 dpi | Shedding | *p* value | Control | 20 dpi | Shedding | *p* value |
| **Left Parietal** | Dorsal | 9.7 ± 2.3 | 5.5 ± 1.2 | 8.0 ± 3.6 | > 0.05 | 6.7 ± 0.6 | 5.0 ± 1.4 | 3.7 ± 1.2 | > 0.05 |
|  | Ventral | 11.3 ± 2.3 | 16.0 ± 4.5 | 13.3 ± 0.6 | > 0.05 | 6.7 ± 0.6 | 7.3 ± 0.5 | 8.7 ± 1.2 | > 0.05 |
| **Visceral** | Dorsal | 10.7 ± 2.5 | 10.8 ± 4.6 | 14.7 ± 4.0 | > 0.05 | 5.3 ± 0.6 | 5.3 ± 0.1 | 7.7 ± 1.5 | > 0.05 |
|  | Ventral | 14.7 ± 3.2 | 16.8 ± 2.1 | 22.0 ± 1.7 | < 0.05 (*) | 6.3 ± 0.6 | 6.0 ± 0.0 | 7.0 ± 1.7 | > 0.05 |

**Table II.** Gray value intensities of Bgl-FaRP1, Bgl-FaRP2 and Bgl-FaNaC in visceral and left parietal ganglia 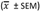

|  |  | *Bgl*-FaRP1 |  |  |  | *Bgl*-FaRP2 |  |  |  | *Bgl*-FaNaC |  |  |  |
| --- | --- | --- | --- | --- | --- | --- | --- | --- | --- | --- | --- | --- | --- |
|  |  | Control | 20 dpi | Shedding | *p* value | Control | 20 dpi | Shedding | *P* value | Control | 20 dpi | Shedding | *p* value |
| **Left Parietal** | Dorsal | 6.01 ± 0.51 | 7.96 ± 3.46 | 5.43 ± 0.73 | > 0.05 | 7.58 ± 0.62 | 8.54 ± 1.54 | 7.41 ± 2.19 | > 0.05 | 29.9 ± 2.72 | 24.07 ± 4.14 | 14.19 ± 1.45 | < 0.05 (*) |
|  | Ventral | 5.78 ± 2.5 | 6.72 ± 1.08 | 6.56 ± 1.31 | > 0.05 | 6.60 ± 1.39 | 10.03 ± 2.41 | 8.82 ± 1.89 | > 0.05 | 35.67 ± 4.45 | 23.07 ± 2.31 | 15.56 ± 1.31 | < 0.05 (*) |
| **Visceral** | Dorsal | 4.29 ± 0.29 | 7.15 ± 3.63 | 5.84 ± 1.49 | > 0.05 | 6.24 ± 2.45 | 7.70 ± 2.41 | 6.23 ± 1.26 | > 0.05 | 24.29 ± 3.39 | 19.15 ± 2.11 | 11.21 ± 2.13 | < 0.05 (*) |
|  | Ventral | 7.57 ± 2.17 | 6.34 ± 1.68 | 7.01 ± 3.27 | > 0.05 | 7.42 ± 0.28 | 7.31 ± 0.22 | 5.89 ± 2.32 | > 0.05 | 37.50 ± 5.17 | 24.89 ± 1.25 | 15.64 ± 1.95 | < 0.05 (*) |

The multiplexing capacity of the HCR system enabled comparison between *Bgl*-FaNaC localization and expression of the FaRP precursors. In agreement with previous immunohistochemical observations (Rolón-Martínez et al., 2021), strong expression of the tetrapeptide *Bgl*-FaRP1 precursor occurred in a single pair of neurons in the buccal ganglia (Fig. 7a). The buccal ganglia were devoid of neurons expressing *Bgl*-FaRP2 (Fig. 7b). While strong labeling of the *Bgl*-FaNaC receptor was also observed in a single pair of buccal neurons (Fig. 7c), multiplexed labeling showed that the receptor expression did not colocalize with its peptide agonist (Fig. 7d).

Colocalization of the *Bgl*-FaRP1 and *Bgl*-FaNaC messages was observed in the left ventral lobe (VL) of the cerebral ganglion, a lateralized CNS region involved in penile control (Fig. 8; Delgado et al. 2012; Rolón-Martínez et al. 2021). While the majority of VL neurons that expressed *Bgl*-FaRP1 did not label for *Bgl*-FaNaC, colocalization did occur in a cell cluster in the anterolateral region of the lobe (Fig, 8 d-f). Colocalization of receptor and agonist expression was also observed in two larger left cerebral ganglion cells that were not within the VL (Fig. 8 a-c, g-i). These observations suggest that the *Bgl*-FaNaC receptor could play a presynaptic or autoreceptor role in the circuits that control male reproductive behavior (see Discussion).

**Fig. 8.**
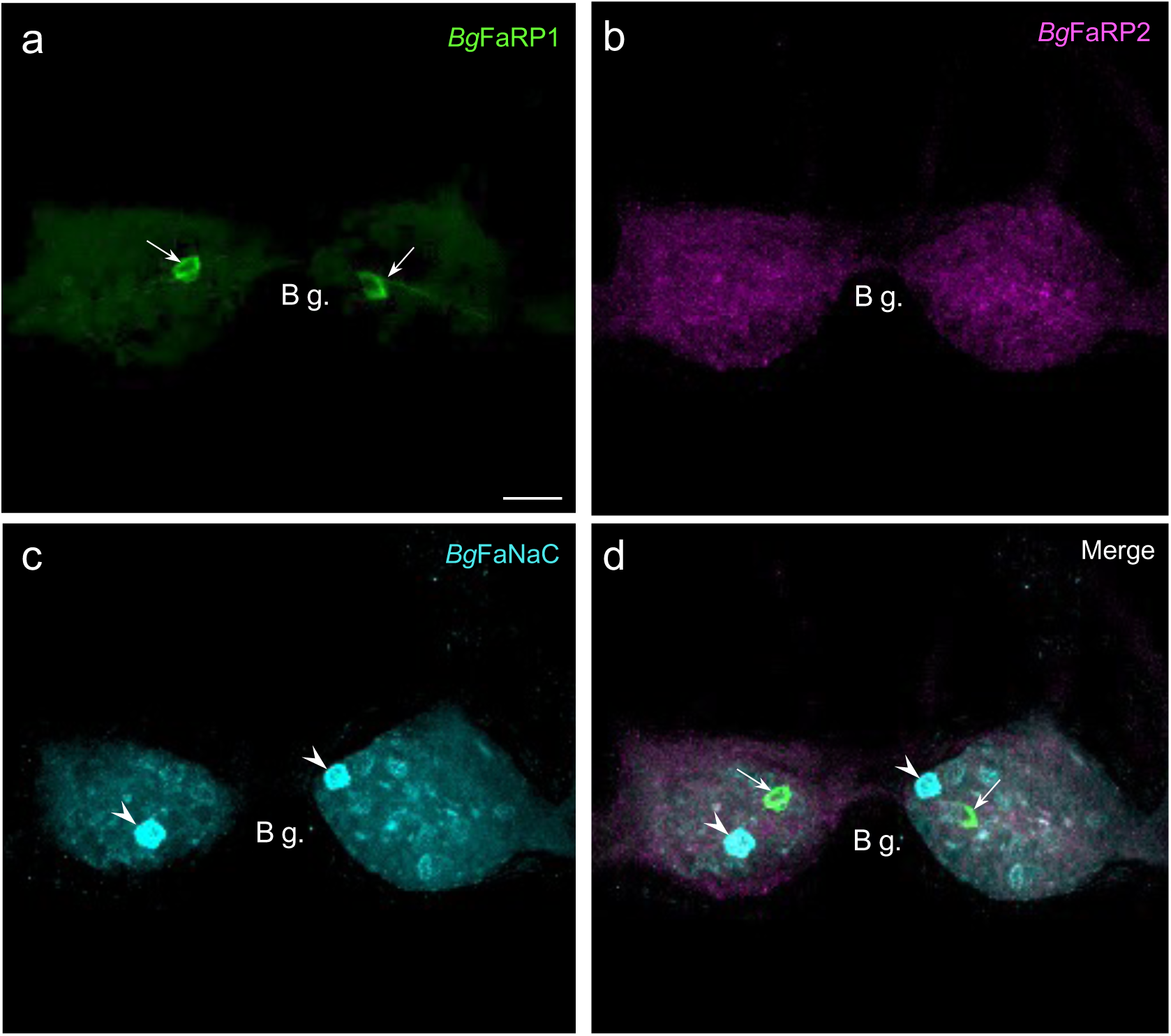
*Bgl*-FaNaC mRNA is not co-expressed with the FMRF-NH_2_ peptide precursors in the buccal ganglion. **a:** Tetrapeptide precursor *Bgl*-FaRP1 mRNA labeling was detected in two neurons in the buccal ganglia *(arrows*, see Rolón-Martínez et al. 2021). *Calibration bar* = 30 µm, applies to all panels. **b:** Expression of the heptapeptide *Bgl*-FaRP2 precursor was not observed in buccal neurons. **c:** Two buccal cells exhibited strong FaNaC mRNA labeling (*arrowheads*). **d:** An overlay of panels a-c showed that the cells expressing FaNaC (*arrowheads*) did not correspond to the neurons expressing FaRP1 (*arrows*).

Localization of *Bgl*-FaRP1 and *Bgl*-FaRP2 expression in the visceral and left parietal ganglia agreed with immunohistochemical findings obtained with precursor specific antibodies (Rolón-Martínez et al. 2021). Abundant *Bgl*-FaRP1 expression was observed in the anterolateral E group (Egp) of cells in the visceral ganglion and the B group (Bgp) of the left parietal ganglion (Fig. 9a). The heptapeptide *Bg*FaRP2 precursor mRNA was expressed in the posterolateral F group (Fgp) of the visceral ganglion and in a posteromedial cluster in the left parietal ganglion (Fig. 9b). *Bgl*-FaNaC was expressed in neurons spanning the region between the Egp and the Fgp on the ventral surface of the visceral ganglion (Fig. 9c). Although a few of the *Bgl*-FaNaC cells overlapped with the E and F groups of the visceral ganglion, no co-expression of the peptide precursors and the receptor was detected (Fig. 9d).

**Fig. 9.**
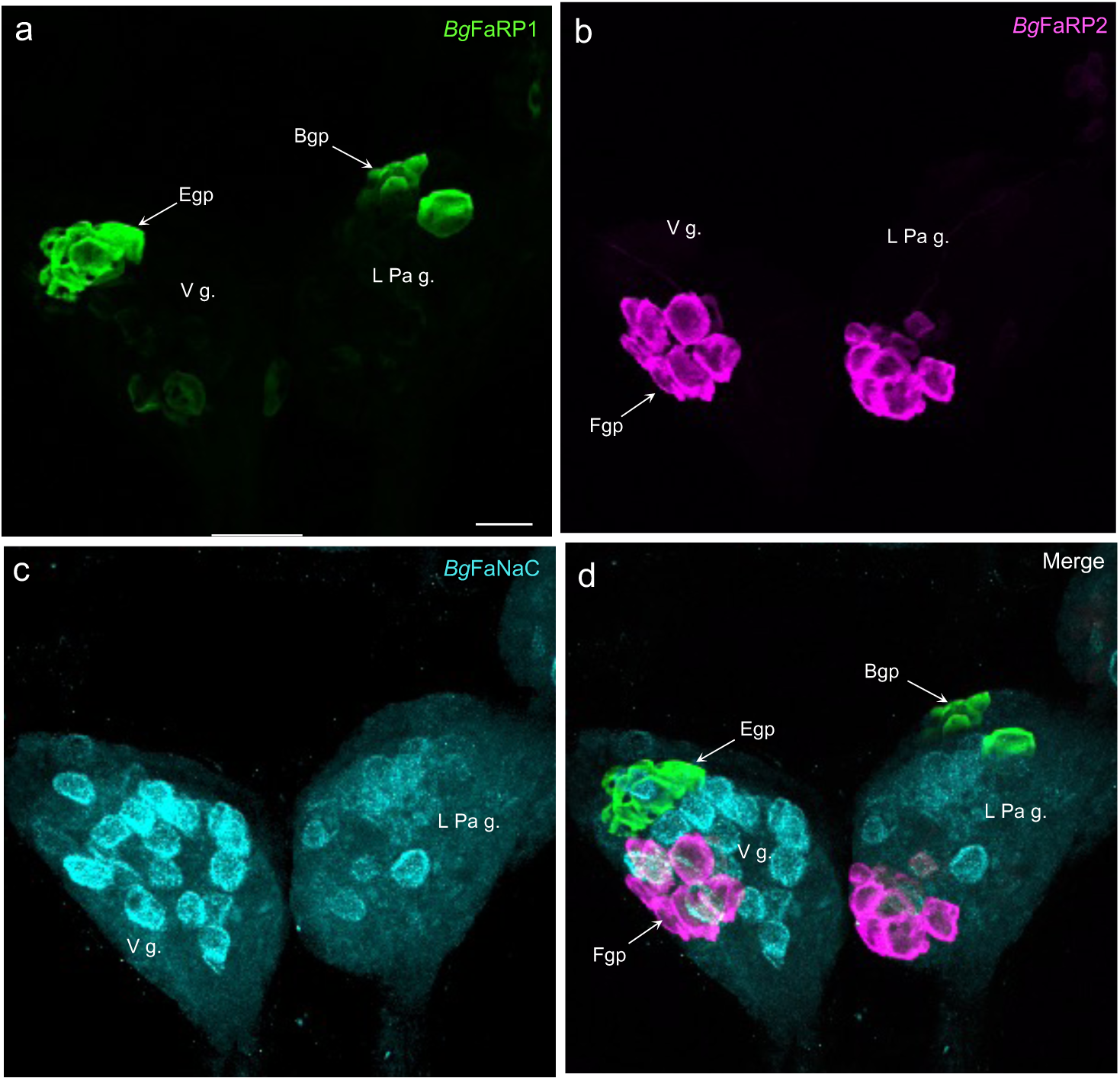
Distinct expression patterns of *Bgl*-FaNaC and the FMRF-NH_2_ precursors in the visceral and left parietal ganglia. **a:** Tetrapeptide precursor *Bgl*-FaRP1 mRNA labeling was detected in the anterolateral E group (*Egp*) of neurons in the visceral ganglion (*V g.*) and in the B group (*Bgp*) of anterolateral cells in the left parietal ganglion (*L Pa g.*). Ventral surface shown. *Calibration bar* = 50 µm applies to all panels. **b:** Expression of the heptapeptide *Bgl-*FaRP2 precursor was observed in the posterolateral F group (*Fgp*) in the visceral ganglion and in a posteromedial cluster of neurons in the left parietal ganglion. **c:** Cells labeled for FaNaC mRNA spanned the central region of the visceral ganglion. Labeling was less intense on the ventral surface of the left parietal ganglion where it also occupied the region between the peptide expressing clusters. **d:** Overlay of panels a-c showed that the cells expressing FaNaC intersected with the tetrapeptide and heptapeptide clusters, but co-expression of the receptor with the peptides was not detected in individual cells.

### 3.3 FMRF-NH_2_ precursor and receptor expression following infection

Due to the multiplexing capability and high resolution attained with the HCR protocol, this approach was utilized for experiments testing potential effects of *S. mansoni* infection on peptide and receptor expression. Nervous systems were dissected from snails that were not exposed to miracidia and size-matched specimens at 20 days post infection (dpi). Infection was verified in a ‘shedding’ group at 35 dpi by confirming release of cercariae upon exposure to light. Expression was quantified by counting the number of cells that displayed an HCR signal at above background intensity levels and by measuring the average intensity of labeled cells. Summary data showed that the number of cells expressing the *Bgl-*FaRP2 heptapeptide precursor was unchanged at the time points examined (Fig. 11a-d, visceral ganglion Fgp shown; control: 6.3 ± 0.6 cells; 20 dpi: 6.0 ± 0.0 cells; shedding: 7.0 ± 1.7 cells; ANOVA: *F*_(2,7)_ = 9.68; *p* = 0.05). Mean gray values of the *Bgl-*FaRP2 HCR signals were also unchanged (control: 7.42 ± 0.28; 20 dpi: 7.31 ± 0.22; shedding: 5.89 ± 2.32; ANOVA: *F*_(2,7)_ = 0.58; *p* = 0.58; Fig. 11e). These findings indicate that expression of the FaRP heptapeptide precursor is not affected by *S. mansoni* infection at the time points tested (Tables I & II).

**Fig. 11.**
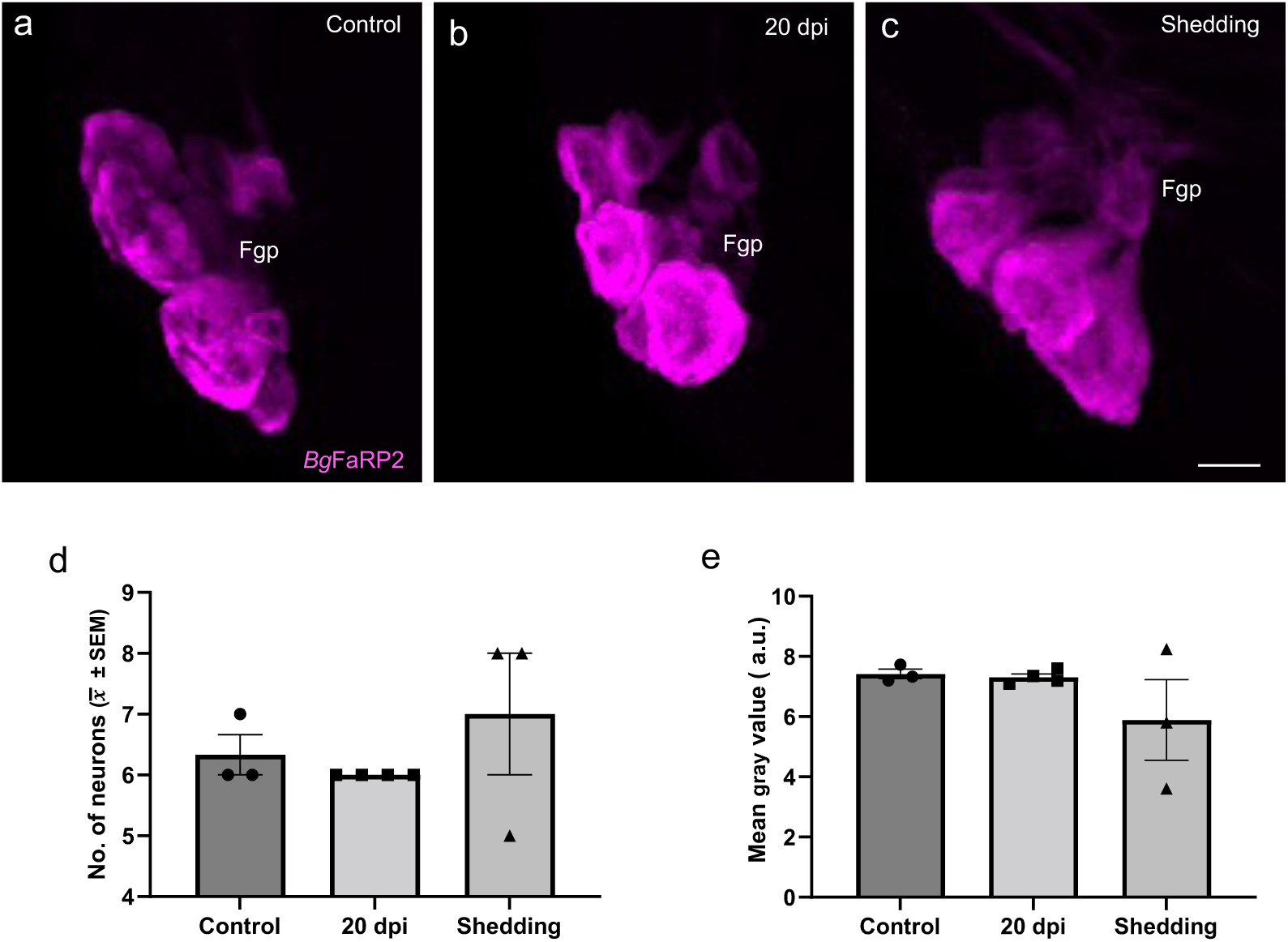
Expression of the heptapeptide *Bgl*-FaRP2 precursor in the visceral ganglion was not altered following *S. mansoni* infection. **a:** In uninfected specimens, FaRP2 expression in the visceral F group was detected in 6.3 ± 0.6 cells with a mean grey value 37.50 ± 5.17. **b:** Differences in expression were not measured at 20 dpi or **c:** in shedding snails. *Calibration bar* = 30 µm. **d:** Summary data verified that the number of visceral Fgp neurons expressing FaRP2 did not differ from control levels at 20 dpi or shedding. **e:** Summary data confirmed that the mean gray value for Fgp neurons expressing FaRP2 did not differ from control levels at 20 dpi or in shedding specimens (a.u.: arbitrary units)..

Unlike the heptapeptide precursor, HCR *in situ* hybridization did disclose an effect of *S. mansoni* infection on the *Bgl*-FaRP1 tetrapeptide precursor (Fig. 12). Effects were analyzed on the Bgp of the left parietal ganglion and the Egp of the visceral ganglion (Table I & II; Fig. 12). Summary data showed that the number of Egp cells expressing the *Bgl-*FaRP1 tetrapeptide precursor increased in the shedding (Fig. 12 a-c, e; control: 14.7 ± 3.2 cells; 20 dpi: 16.8 ± 2.1 cells; shedding: 22.0 ± 1.7 cells; ANOVA: *F*_(2,7)_ = 7.29; *p* = 0.03). Mean gray values of the Egp *Bgl*-FaRP1 HCR signals were not significantly changed (control: 7.57 ± 2.17; 20 dpi: 6.34 ± 1.68; shedding: 7.01 ± 3.27; ANOVA: *F*_(2,7)_ = 0.21; *p* = 0.82; Fig. 12d). Together, these observations indicate that an increased number of neurons within the Egp express high levels of the tetrapeptide (FMRF-NH_2_) precursor in infected snails. Overall levels of expression, per cell, do not appear to be changed.

**Fig. 12.**
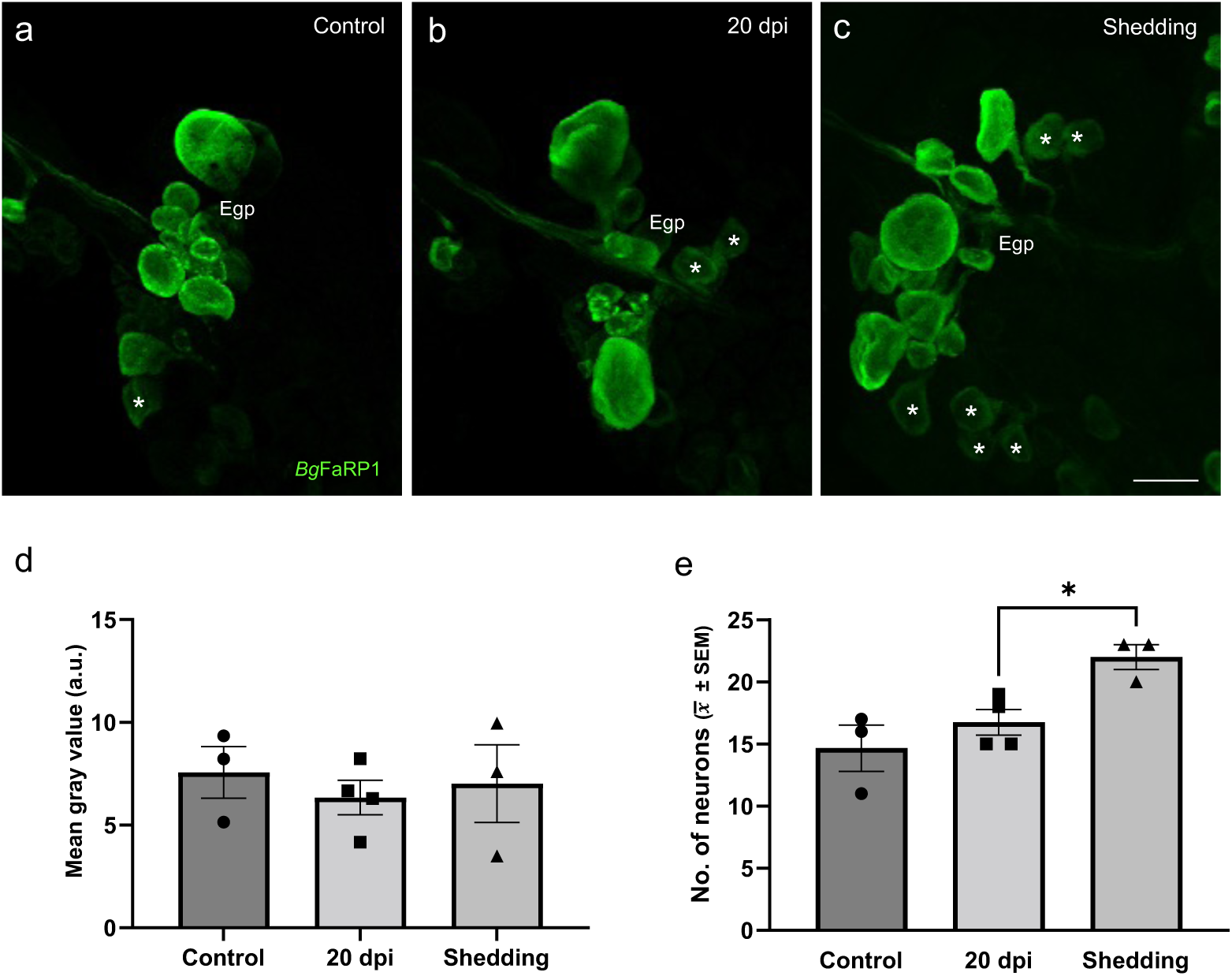
Expression of the tetrapeptide Bgl-FaRP1 precursor in the visceral ganglion was increased following *S. mansoni* infection. **a:** In uninfected specimens, FaRP1 expression in the visceral E group was detected in 14.7 ± 3.2 cells with a mean grey value 7.57 ± 2.17 (Tables I & 2). **b:** Differences in expression were not measured at 20 dpi. **c:** In shedding snails, the number of Egp neurons expressing FaRP1 was increased. *Calibration bar* = 30 µm applies to all panels. **d:** Summary data verified that the mean gray value of visceral Egp neurons expressing FaRP1 did not differ from control levels at 20 dpi or in shedding specimens. **e:** Summary data confirmed that the number of Egp neurons expressing FaRP1 was increased in shedding specimens. Asterisks indicate cells in which low levels of FaRP1 were detected.

As *Bgl-*FaNaC was not expressed in discrete clusters, changes were assessed with gray values measured over the visceral and left parietal ganglia surfaces (Table II, Figs. 13, 14). Expression was decreased on the ventral surface of the visceral ganglion (control: 37.50 ± 5.17; 20 dpi: 24.89 ± 1.25; shedding: 15.64 ± 1.95; ANOVA: *F*_(2,7)_ = 32.47; *p* = 0.01; Fig. 13a-d). *Bgl*-FaNaC mRNA labeling was also decreased on the dorsal surface of the visceral ganglion of infected snails (control: 24.29 ± 3.39; 20 dpi: 19.15 ± 2.11; shedding: 11.21 ± 2.13; ANOVA: *F*_(2,7)_ = 18.80; *p* = 0.01; Fig. 13e-h). While there was a tendency toward lower expression at 20 dpi on both surfaces, the decreases only reached significant values in shedding specimens.

**Fig. 13.**
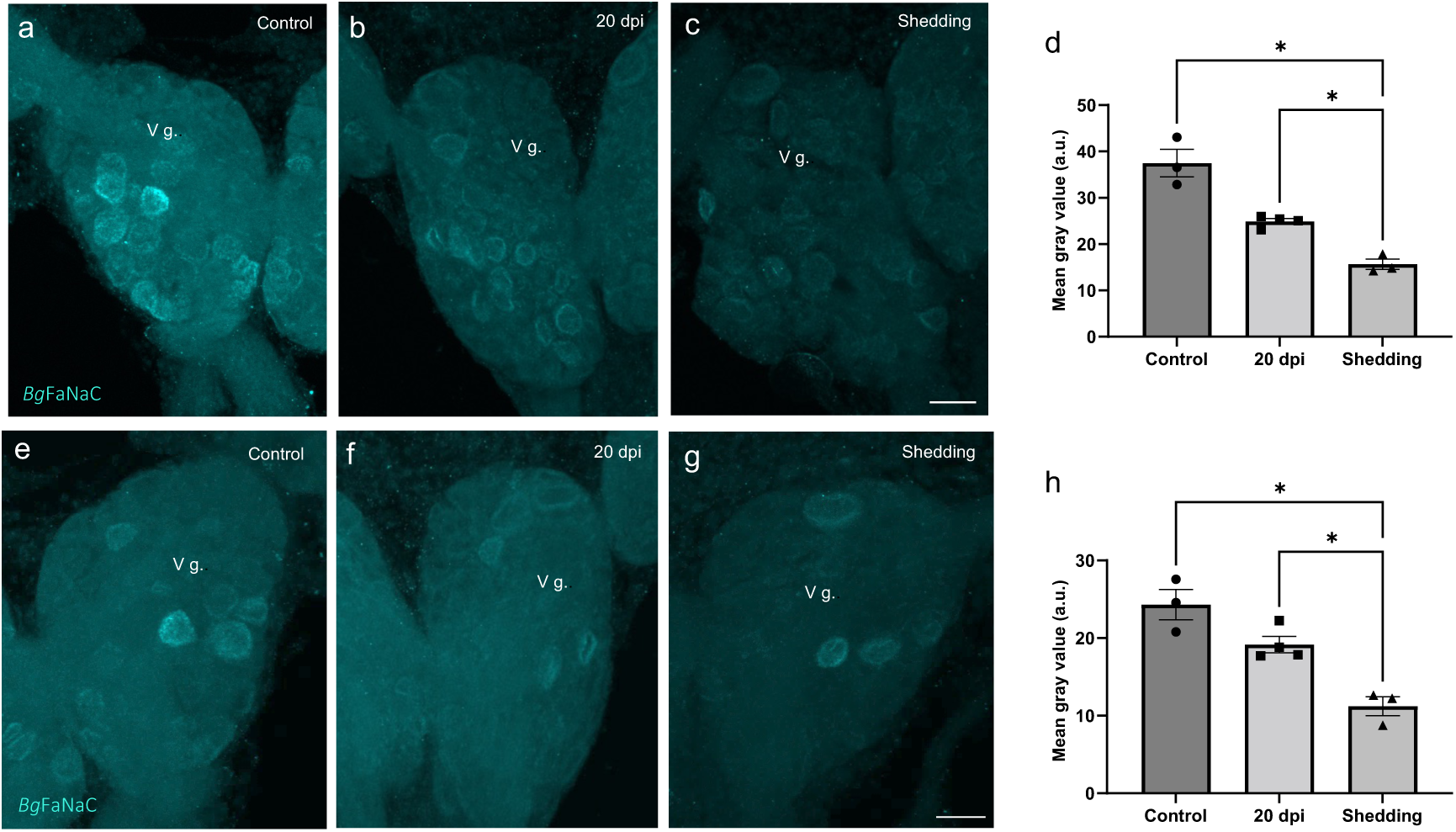
Expression of the FaNaC receptor in the visceral ganglion was decreased following *S. mansoni* infection. **a:** In uninfected specimens, FaNaC expression produced a mean grey value of 37.50 ± 5.17 on the ventral surface of the visceral ganglion (Table II). **b, c:** Decreased gray values were measured at expression was observed at 20 dpi and in shedding snails. *Calibration bar* = 30 µm applies to a-c. **d:** Summary data verified that the mean gray value of FaNaC expression in ventral visceral neurons decreased across the chronology of infection. **e:** In uninfected specimens, FaNaC expression produced a mean grey value 24.29 ± 3.39 on the dorsal surface of the visceral ganglion. **f, g:** Decreased expression was observed at 20 dpi and in shedding snails. *Calibration bar* = 30 µm applies to e-g. **h:** Summary data verified that mean gray value of FaNaC expression in dorsal visceral neurons decreased across the chronology of infection.

**Fig. 14.**
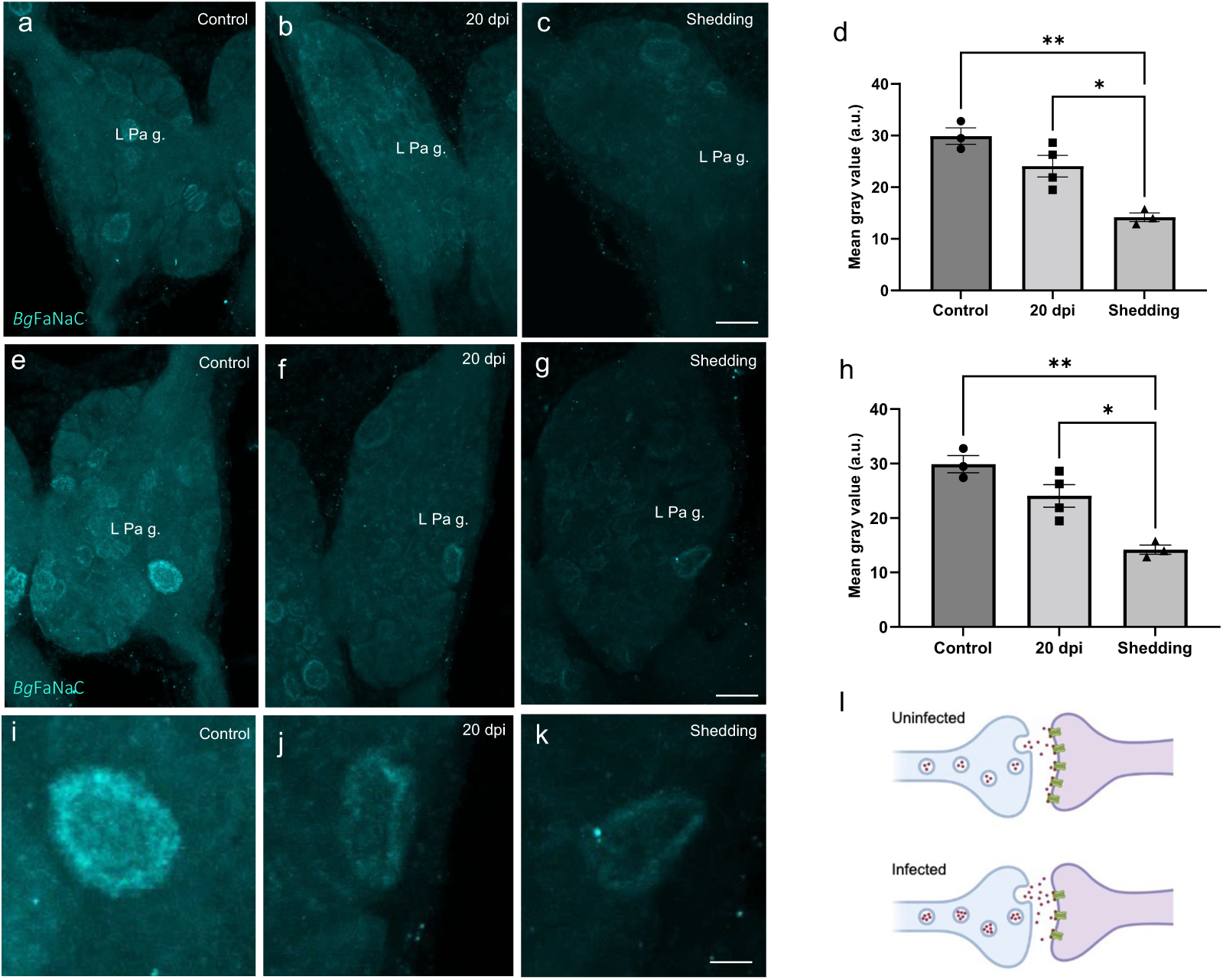
Expression of the *Bgl*-FaNaC receptor in the left parietal ganglion was decreased following *S. mansoni* infection. **a:** In uninfected specimens, a mean grey value of 29.90 ± 2.72 FaNaC expression was measured on the dorsal surface of the left parietal ganglion. **b:** Decreased expression was observed at 20 dpi. **c:** In shedding snails, the number of dorsal left parietal neurons expressing FaNaC and the mean level of expression were both decreased. *Calibration bar* = 30 µm applies to a-c. **d:** Summary data confirmed that the mean gray value of FaNaC expression in dorsal left parietal neurons decreased across the chronology of infection. **e:** In uninfected specimens, a mean gray value of 35.67 ± 4.45 FaNaC expression was measured on the ventral surface of the left parietal ganglion. **f:** Decreased expression was observed at 20 dpi. **g:** In shedding snails, the number of ventral left parietal neurons expressing FaNaC and the mean level of expression were both decreased. *Calibration bar* = 30 µm applies to e-g. **h:** Summary data verified that the mean gray value of FaNaC expression in ventral left parietal neurons decreased across the chronology of infection. **i-k:** Large ventral left parietal ganglion neuron (panels e-g) shown at higher magnification. **l:** Schematic summary of proposed compensatory effects on FMRF-NH2signaling followingS. *mansoni* infection. While expression of the peptide agonist (red circles) is increased following infection, expression of the FaNaC postsynaptic channels (green) is decreased. Such opposing actions could exert a stabilizing influence on FMRF-NH2ligand-gated signaling.

*Bgl*-FaNaC expression was also decreased in the left parietal ganglion of infected snails (Table II). On the dorsal surface, decreased expression reached significance at the shedding stage (control: 29.90 ± 2.72; 20 dpi: 24.07 ± 4.14; shedding: 14.19 ± 1.45; ANOVA: *F*_(2,7)_ = 22.5; *p* = 0.002; Fig. 14a-d). *Bg*FaNaC expression was also decreased on the ventral surface of the left parietal ganglion (control: 35.67 ± 4.45; 20 dpi: 23.07 ± 2.31; shedding: 15.56 ± 1.31; ANOVA: *F*_(2,7)_ = 33.93; *p* = 0.006; Fig. 14e-k). The decrease on the ventral surface reached significant levels at 20 dpi (Fig. 14h).

## 4 Discussion

### 4.1 Properties of the *Bgl*-FaNaC

The FaNaC receptors that have been studied to date exhibit at least a 35-fold range of efficacy, with EC_50_ values for FMRF-NH_2_ varying from 2 × 10^−6^ M in *Cornu aspersum* (Lingueglia et al. 1995) to 7 × 10^−5^ M in *Planorbella trivolvis* (Jeziorski et al. 2000). The EC_50_ of 3.3 × 10^−4^ M observed for the *Biomphalaria* FaNaC in the present investigation may be considered in the context of structure-activity studies on the *Cornu* and *Planorbella* receptors. Chimeras constructed from these receptors indicated that the peptide recognition site is located in the extracellular region following TM1 (shaded red in Fig. 1; Cottrell et al. 2001; Cottrell 2005). Within this region, site-directed mutagenesis was used to substitute *Planorbella* amino acids for the *Cornu aspersum* residues at positions Y131Q, N134T, or I160F (Niu et al. 2016; enclosed by rectangles in Fig. 1). Each substitution produced a FMRF-NH_2_ EC_50_ value that was significantly higher than the native *C. aspersum* receptor (Niu et al. 2016). Notably, these three ‘low affinity’ residues are conserved between the planorbids *Planorbella* and *Biomphalaria*. A fourth substitution that also significantly reduced the affinity of FMRF-NH_2_, D154K (dashed rectangle, Fig. 1), is not shared between *Planorbella* and *Biomphalaria* and may contribute to the apparent 4-fold difference in their efficacy.

Divergent peptide recognition sequences could also account for species differences in agonist specificity. FLRF-NH_2_, which is present in two copies on the *B. glabrata* FMRF-NH_2_ precursor (Rolón-Martínez et al. 2021), activates the *C. aspersum* FaNaC with an EC_50_ of 11 µM (Lingueglia et al. 1995). In contrast, responses of the *Planorbella* FaNaC to FLRF-NH_2_ were negligible (Jeziorski et al. 2000), in agreement with our observations on *Biomphalaria*. The high level of sequence conservation between the planorbids *Cornu* and *Biomphalaria* (>90%) may therefore confer an extraordinary degree of agonist specificity in addition to the reduced efficacy of FMRF-NH_2_. The possibility that FLRF-NH_2_ could act as an antagonist at the *Bgl*-FaNaC was not tested.

### 4.2 Localization of the *Bg*FaNaC

Our findings that *Bgl*-FaNaC mRNA was confined to cell bodies and its abundant expression in the subesophageal (visceral and parietal) ganglia were consistent with observations in *Planorbella trivolvis* and *Helix aspersa* (Davey et al. 2001). The predominant localization of the *Bgl*-FaNaC protein to neuronal processes was also in agreement with immunohistochemical observations in *Planorbella trivolvis*. When FMRF-NH_2_ was applied to isolated giant dopaminergic neurons (GDN; corresponding to LPeD1 of *Biomphalaria*; Fig. 6a, b) large inward currents were produced with focal application near the axon hillock, leading to the suggestion that newly synthesized membrane channels were inserted at a high density prior to translocation to distal sites (Davey et al. 2001).

It is well established that the FMRF-NH_2_ related peptides participate in multiple neural circuits in gastropods, including the control of feeding motor programs (Sossin et al. 1987; Murphy 1990; Alania et al. 2004), male mating behavior (Van Golen et al. 1995; de Lange et al. 1998a b), and cardiorespiratory regulation (Buckett et al. 1990; Worster et al. 1998). *Bgl*-FaNaC involvement in each of these circuits was supported by its expression patterns in the buccal (Fig. 7), cerebral (Fig. 8), and visceral (Fig. 9) ganglia, respectively. In the feeding and cardiovascular networks, *Bgl*-FaNaC was expressed in neurons that were in close proximity to cells that express the *Bgl*-FaRP1 precursor.

Co-expression of the *Bgl*-FaNaC and the message for the FMRF-NH_2_ tetrapeptide precursor was rare, but instances were detected in regions of the cerebral ganglion that control male mating behavior (Fig. 8). Such co-expression of an agonist and its receptor could enable neurons to form autapses (see Van der Loos and Glaser 1972). Excitatory autaptic signaling has been shown to produce long-lasting afterdischarges in gastropod feeding and reproductive systems (Brussaard et al., 1990; Norekian 1999; Saada et al. 2009). Autapses are proposed to provide a mechanism whereby a brief stimulus can produce a prolonged response required to drive a motor circuit. In gastropods, male copulation consists of a stereotyped sequence of actions, including preputium eversion, probing, penis eversion, and intromission (de Lange et al. 1998a, b; Koene 2010). Each action lasts for several minutes, probably persisting after termination of its initiating stimulus. Interestingly, application of FMRF-NH_2_ to the water surrounding *Biomphalaria* caused preputium eversion, a behavior that lasts several minutes in the copulatory sequence (Fong et al. 2005). The larger cerebral neurons in which *Bgl*-FaNaC and *Bgl*-FaRP1 are co-expressed (Fig. 8g-i) could provide opportunities to examine the involvement of FaNaC receptors in autapses.

### 4.3 Response to infection

Due to the pleiotropic functions of the FaRPs in gastropods, this neuropeptide signaling system is considered a potential target for schistosome larvae (de Jong-Brink 1995; de Jong-Brink et al. 2001). The increase in *Bgl*-FaRP1 expression observed this study (Fig. 12, Table I) agrees with previous studies that measured neuropeptide responses to infection in gastropods. In the host-parasite interaction between the pulmonate snail *Lymnaea stagnalis* and the avian schistosome *Trichobilharzia ocellata*, significant increases in FMRF-NH_2_ gene expression were measured across the post-infection chronology (Hoek et al. 1997). The early onset of this increase (>300% at five hours) was suggested to reflect a direct effect of parasitism on the host brain. A lower (<100%) increase at later time points (6 and 8 weeks post-infection) was proposed to contribute to the schistosome survival strategy during the shedding stage of infection, when host energy resources are redirected toward the large numbers of cercariae inhabiting the snail. Elevated levels of FMRF-NH_2_ were also detected in *B. glabrata* at 12 days post-infection with *S. mansoni* (Wang et al. 2017). Of 39 CNS peptides that exhibited >1.5-fold changes, FMRF-NH_2_ was one of only 6 that was increased. It was proposed that the increased expression of FMRF-NH_2_ could contribute to enhanced metabolic activity during the pre-patent phase of infection (Wang et al. 2017).

Our observations suggest one mechanism that could contribute to the elevated levels of FMRF-NH_2_ in infected snails (Fig. 14l). Increased precursor expression was limited to *Bgl*-FaRP1, the tetrapeptide precursor that encodes FMRF-NH_2_, the sole *Bgl*-FaNaC agonist (Fig. 12, Table I). No changes in expression were observed for the heptapeptide precursor *Bgl*-FaRP2 (Fig. 11, Tables I & II). Moreover, the increased expression was limited to a subset of *Bgl*-FaRP1expressing neurons in the visceral ganglion and was primarily observed late in the infection chronology. In contrast, down-regulation of the *Bgl*-FaNaC receptor appeared to commence earlier and occurred throughout the visceral and left parietal ganglia. We propose that the increased *Bgl*-FaRP1 expression could reflect a compensatory mechanism that occurs in response to decreased receptor expression (Fig. 14l). Such homeostatic increases in neuropeptide expression would only occur in neurons that are presynaptic to neurons that express *Bgl*-FaNaC. Future studies should explore the role of *Bgl*-FaNaC in synaptic signaling and examine whether such signaling is maintained despite reduced receptor expression levels following infection. The limited phylogenetic scope of this signaling pathway, and its apparent involvement in multiple vital physiological and behavioral circuits, could provide novel strategies for control of snail pests.

## Acknowledgments

Supported by the National Institutes of Health: MD007600 (RCMI), GM103642 (COBRE), GM007821 (MARC), GM061838 (RISE); National Science Foundation: DBI-0932955, HRD-1137725, OISE-1545803, and DBI-1337284; National Academy of Sciences (NAS; USA) U.S.-Egypt Science and Technology (S&T) Joint Fund 2000007152*; Science and Technology Development Fund (STDF, Egypt). The authors thank Dr. Margaret Mentink-Kane and the staff at the Biomedical Research Institute (BRI) NIH-NIAID Schistosomiasis Resource Center for providing infected specimens.

